# Climate change will drive novel cross-species viral transmission

**DOI:** 10.1101/2020.01.24.918755

**Authors:** Colin J. Carlson, Gregory F. Albery, Cory Merow, Christopher H. Trisos, Casey M. Zipfel, Evan A. Eskew, Kevin J. Olival, Noam Ross, Shweta Bansal

## Abstract

At least 10,000 species of mammal virus are estimated to have the potential to spread in human populations, but the vast majority are currently circulating in wildlife, largely undescribed and undetected by disease outbreak surveillance ^1–3^ . In addition, changing climate and land use are already driving geographic range shifts in wildlife, producing novel species assemblages and opportunities for viral sharing between previously isolated species ^4, 5^ . In some cases, this will inevitably facilitate spillover into humans ^6, 7^ —a possible mechanistic link between global environmental change and emerging zoonotic disease ^8^ . Here, we map potential hotspots of viral sharing, using a phylogeographic model of the mammal-virus network, and projections of potential geographic range shifts for 3,139 mammal species under climate change and land use scenarios for the year 2070. Range-shifting mammal species are predicted to aggregate at high elevations, in biodiversity hotspots, and in areas of high human population density in Asia and Africa, driving the novel cross-species transmission of their viruses an estimated 4,000 times. Counter to expectations, holding warming under 2°C within the century does not reduce new viral sharing, due to greater potential range expansions— highlighting the need to invest in surveillance even in a low-warming future. Most projected viral sharing is driven by diverse hyperreservoirs (rodents and bats) and large-bodied predators (carnivores). Because of their unique dispersal capacity, bats account for the majority of novel viral sharing, and are likely to share viruses along evolutionary pathways that could facilitate future emergence in humans. Our findings highlight the urgent need to pair viral surveillance and discovery efforts with biodiversity surveys tracking species’ range shifts, especially in tropical countries that harbor the most emerging zoonoses.

## Main Text

In the face of rapid environmental change, survival for many species depends on moving to track shifting climates. Even in a best case scenario, many species’ geographic ranges are projected to shift a hundred kilometers or more in the next century ^9, 10^. In the process, many animals will bring their parasites and pathogens into new environments ^4, 11^. This poses a measurable threat to global health, particularly given several recent epidemics and pandemics of viruses that originate in wildlife (zoonotic viruses, or zoonoses) ^1, 12, 13^. Most frameworks for predicting cross-species transmission therefore focus on the steps that allow animal pathogens to make the leap to human hosts (a process called spillover) ^13–15^. However, zoonotic viruses are a small fraction of total viral diversity, and viral evolution is an undirected process ^16^, in which humans are only one of over 5,000 mammal hosts with over 12 million possible pairwise combinations ^17^ (to say nothing of the other four classes of vertebrates, which have a much greater fraction of undescribed viral diversity). If those host species track shifting climates, they will share viruses not just with humans, but with each other, for the very first time ^8^. Despite their indisputable significance, spillover events are probably just the tip of the iceberg; by numbers alone, most cross-species transmission events attributable to climate change will probably occur among wildlife hosts, potentially threatening wildlife populations and largely undetected by zoonotic disease surveillance.

The scale of this process will depend on *opportunity* and *compatibility* ^13, 18, 19^, and both dimen-sions pose an important predictive challenge. Because only a few species are common world-wide, most hosts have no *opportunity* to exchange pathogens: of all possible pairs of mammal species, only *∼*7% share any geographic range, and only *∼*6% are currently known to host one or more of the same virus species (hereafter *viral sharing*) ^18^. As host geographic ranges shift, some interactions will become possible for the first time, and a subset will lead to viral establishment in a previously-inaccessible host (*novel viral sharing*). The potential ability of species to track shifting climate and habitat conditions will determine which pairs of species encounter each other for the first time ^4, 20^. Even if species’ ranges nominally overlap, habitat selection and behavioral differences can further limit contact ^20^. Although some viruses spread environmentally or by arthropod vectors between spatially proximate species with no direct behavioral contact ^21^, sharing is more likely on average among species with more ecological overlap ^22^. Even among species in close contact, most cross-species transmission events are still a dead end. Progressively smaller subsets of viruses can infect novel host cells, proliferate, cause disease, and transmit onward in a new host ^19^. Their ability to do so is determined by *compatibility* between viral structures, host cell receptors, and host immunity ^6^. Because closely-related species share both ecological and immunological traits through identity by descent, phylogeny is a strong predictor of pathogen sharing ^18, 23^ and of susceptibility to invasion by new viruses ^24–26^. In a changing world, these mechanisms can help predict how ecosystem turnover could impact the global virome.

Although several studies have mapped current hotspots of emerging diseases ^3, 12, 27^, few have forecasted them in the context of global change. With the global reassortment of animal biodiversity due to climate and land use change, it is unknown whether bats and rodents will still play a central role in viral emergence ^3, 28^ (ED Figure 1), or whether hotspots of vi-ral emergence will stay in tropical rainforests ^27, 29^, which currently harbor most undiscovered viruses ^3, 30^. Here, by projecting potential geographic range shifts (that is, newly suitable habitat, which a species may or may not migrate to) and applying mechanistic biological rules for crossspecies transmission, we predicted how and where global change could potentially create novel opportunities for viral sharing, with particular attention to the potential connections between these risks and human health. We focused on mammals because they have some of the most complete biodiversity data, the highest proportion of viral diversity described ^1^, and the greatest downstream relevance to human health and zoonotic disease emergence of any vertebrate class. We built species distribution models (SDMs) for 3,870 placental mammal species, and projected potential geographic range shifts based on four paired scenarios for climate change (Representative Concentration Pathways, RCPs) and land use change (Shared Socioeconomic Pathways, SSPs) by 2070. These scenarios characterize alternative futures for global environmental change, from sustainable land use change and a high chance of keeping global warming under 2*^◦^* C (SSP1-RCP2.6), to a high chance of 4*^◦^* C warming, continued fossil fuel reliance, and rapid land degradation and change (SSP5-RCP8.5; see “Methods” for a detailed explanation). We present results for SSP1-RCP 2.6 in the main text because this scenario is most in line with the goals of the Paris Agreement to keep global warming “well below” 2*^◦^* C ^31^. We quantified model uncertainty in projected climate futures using nine global climate models (GCM) from the Coupled Model Intercomparison Project Phase 6 (CMIP6). Because many species are unlikely to be biologically suited for rapid range shifts, and will therefore move slower than the local velocity of climate change, we constrained the speed of range shifts based on inferred allometric scaling of animal movement ^32^, and compared scenarios that assumed limited dispersal against “full dispersal” (that is, no dispersal limitation).

We used projections of newly suitable habitat to identify where novel range overlap among currently non-overlapping species could happen (hereafter *first encounters*). We then used a recently-developed viral sharing model to predict the probability of a *novel viral sharing event*— here defined as the future cross-species transmission of at least one virus species, in this case between a pair of hosts that are newly in contact—based on novel geographic overlap and host phylogenetic similarity ^18^, a first order approximation of opportunity and compatibility (ED Figure 2). This model framework has previously provided insights into viral macroecology and zoonotic risk based on the *∼*1% of the global mammalian virome that has been described ^1, 3, 18^.

Based on the total number and distribution of first encounters among a subset of 3,139 species (see “Methods”), we used cumulative viral sharing probabilities to estimate the total number of novel sharing events that are expected (each of which describes the cross-species transmission of at least one virus). Using this approach, we tested the hypothesis that environmental change should alter mammal communities in ways that expose hosts to novel viruses, altering the structure of the whole mammal-virus network.

### Climate and land use change will transform the global virome

If species range shifts can keep pace with the velocity of climate change (i.e., can disperse to all newly suitable locations) ^33^, we predict that the vast majority of mammal species will overlap with at least one unfamiliar species somewhere in their potential future range, regardless of emissions scenario (mean across GCMs *±* s.d. here and after; RCP 2.6: 98.6% *±* 0.2%; RCP 8.5: 96.6% *±* 0.8%). At the global level, geographic range shifts would permit over 300,000 first encounters in every climate scenario (SSP1-RCP 2.6: 316,426 *±* 1,719; SSP5-RCP 8.5: 313,973 *±* 2,094; ED Figure 3). Compared to a present-day baseline, in which we calculated 345,850 current pairwise overlaps among the 3,870 species (*∼* 7%), this essentially represents a doubling of potential species contact. These “first encounters” between mammal species will occur everywhere in the world, but are concentrated in tropical Africa and southeast Asia (ED Figure 4). This result was counter to expectations that species might aggregate at higher latitudes, given that most research has focused on poleward range shifts ^34–36^, and previous work has anticipated a link between climate change, range shifts, and parasite host-switching in the Arctic ^37, 38^. However, we find that when species shift along latitudinal gradients, they travel in the same direction as others that are already included in their assemblage, leading to few first encounters. In contrast, when species track thermal optima along elevational gradients (allowing them to come from different directions; i.e., mountains force species to cluster), they will aggregate in the most novel combinations in mountain ranges, especially in tropical areas with the highest baseline diversity, matching prior predictions ^39^. This pattern was robust to climate model uncertainty (Supplemental Figures 1-9) and to differences in dispersal capacity (e.g., Figure 2C). The most notable model variation is in the Amazon basin, as well as a small portion of the central African basin, Botswana, and parts of the Indian subcontinent (ED Figure 5). These areas become essentially devoid of first encounters in the most sensitive climate models and warmest pathways, presumably because all are high-endemism basins of homogenous climate that may warm too much for species to “escape” into high-elevation refugia (a fairly well-documented pattern ^40–42^).

**Figure 1:**
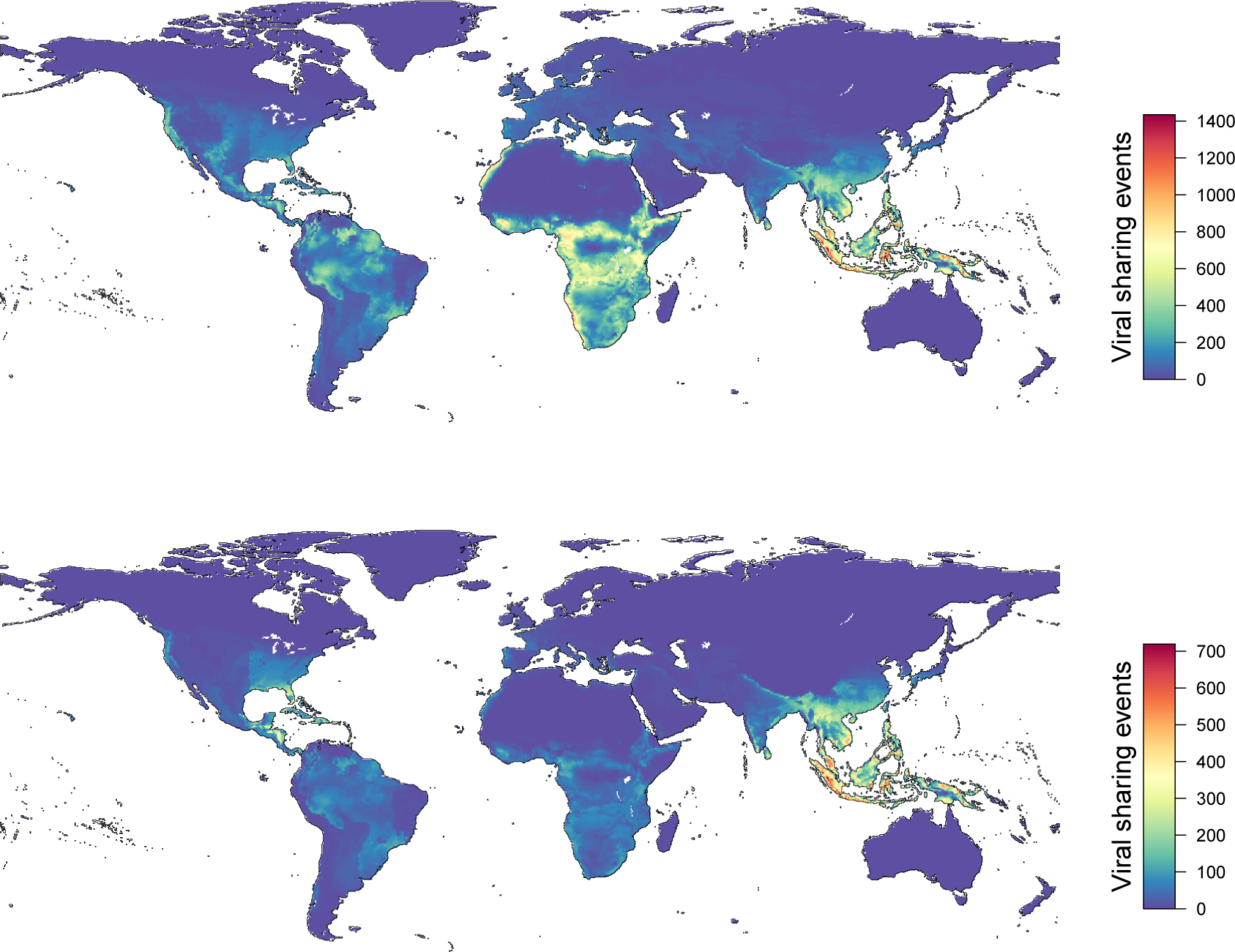
Climate change will drive novel viral sharing among mammal species. The projected number of novel viral sharing events among mammal species in 2070 based on host species geographic range shifts from climate and land use change (SSP1-RCP 2.6), without dispersal limits (A) and with dispersal limitation (B). Results are averaged across nine global climate models.

**Figure 2:**
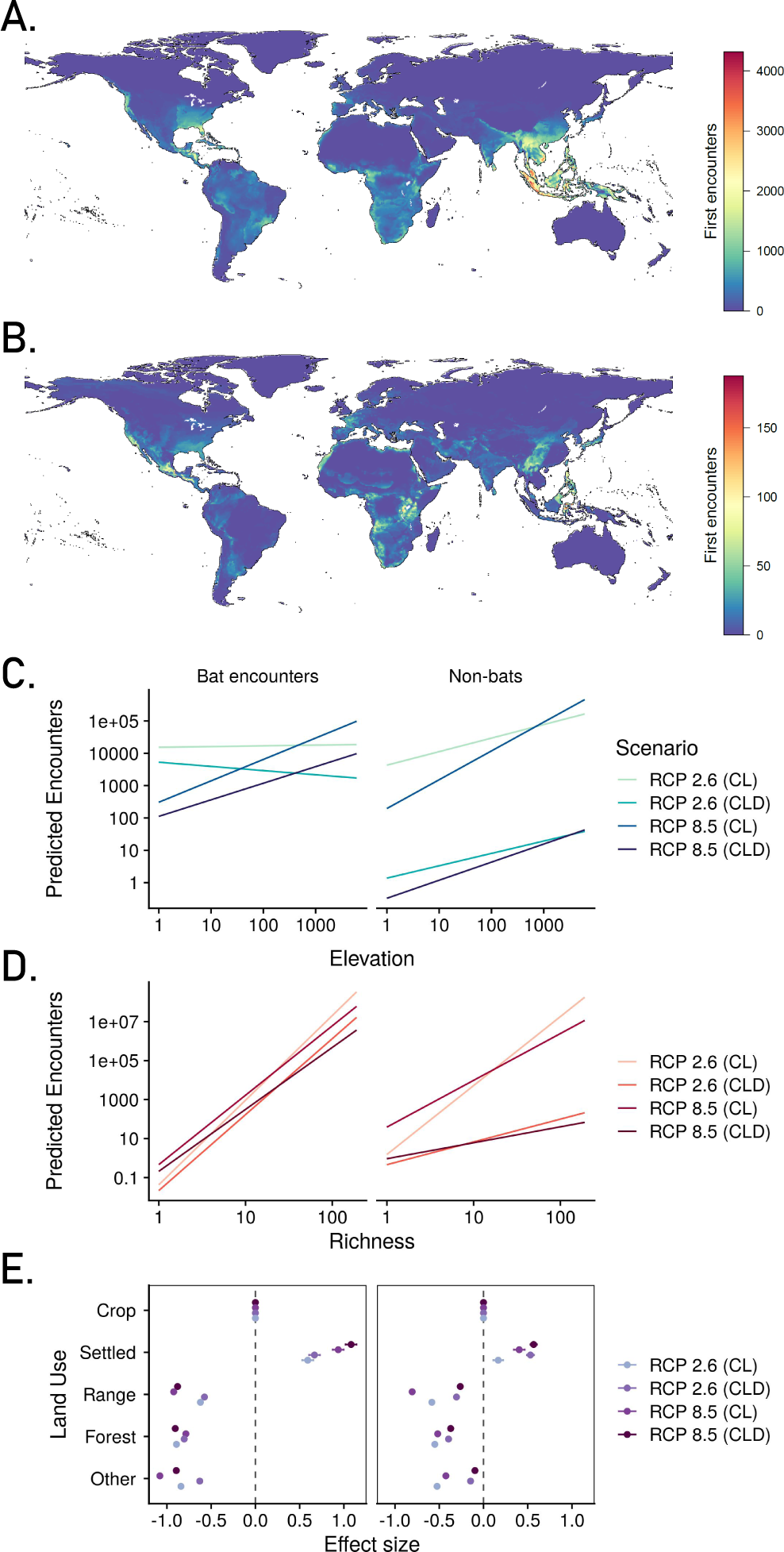
Bats disproportionately drive future novel viral sharing. The spatial pattern of first encounters (in SSP1-RCP 2.6) differs among range-shifting mammal pairs including bat-bat and bat-nonbat encounters (A) and only encounters among non-bats (B). Using a linear model, we show that elevation (C), species richness (D), and land use (E) influence the number of new overlaps for bats and non-bats across scenarios (RCPs paired with SSPs as described in Methods). Slopes for the elevation effect were generally steeply positive: a log_10_-increase in elevation was associated with between a 0.4-1.41 log_10_-increase in first encounters. Results are averaged across nine global climate models. Legends refer to scenarios: CL gives climate and land use change, while CLD adds dispersal limits.

This global re-organization of mammal assemblages is projected to dramatically impact the structure of the mammalian virome. Accounting for geographic opportunity and phylogenetic compatibility, we project that a total of 316,426 (*±* 1,719) first encounters in RCP 2.6 would lead to 15,311 novel sharing events (*±* 140)—that is, a minimum of at least *∼*15,000 cross-species transmission events of at least one novel virus (but potentially many more) between a pair of naive host species. Assuming that viral sharing will initially be localized to areas of novel host overlap, we mapped expected viral sharing events, and found again that most sharing should occur in high-elevation, species-rich ecosystems in Africa and Asia (Figure 1A). If species survive a changing climate by aggregating in high elevation refugia, this suggests emerging viruses may be an increasing problem for their conservation ^43, 44^. Across scenarios, the spatial pattern of expected sharing events was nearly identical, and was dominated more by the extent of potential range shifts than by underlying community phylogenetic structure (ED Figure 6; Supplemental Figures 10-18). Though previous work has suggested that the phylogenetic structure of mammal communities might drive spatial hotspots of pathogen sharing and emergence ^45^, in our framework, opportunity drives spatial patterns more than compatibility. Given that phylogeny is a strong determinant of viral sharing in the underlying model, this difference from previous studies can probably be explained by evolutionary scale, where prior work focused on primates, and our study includes all mammals. At this broader scale, predicted viral sharing patterns mostly track total richness (see Figure 3b in ^18^), and at finer scales, phylogeny has a stronger effect (see Extended Data Figure 8 for an example).

**Figure 3:**
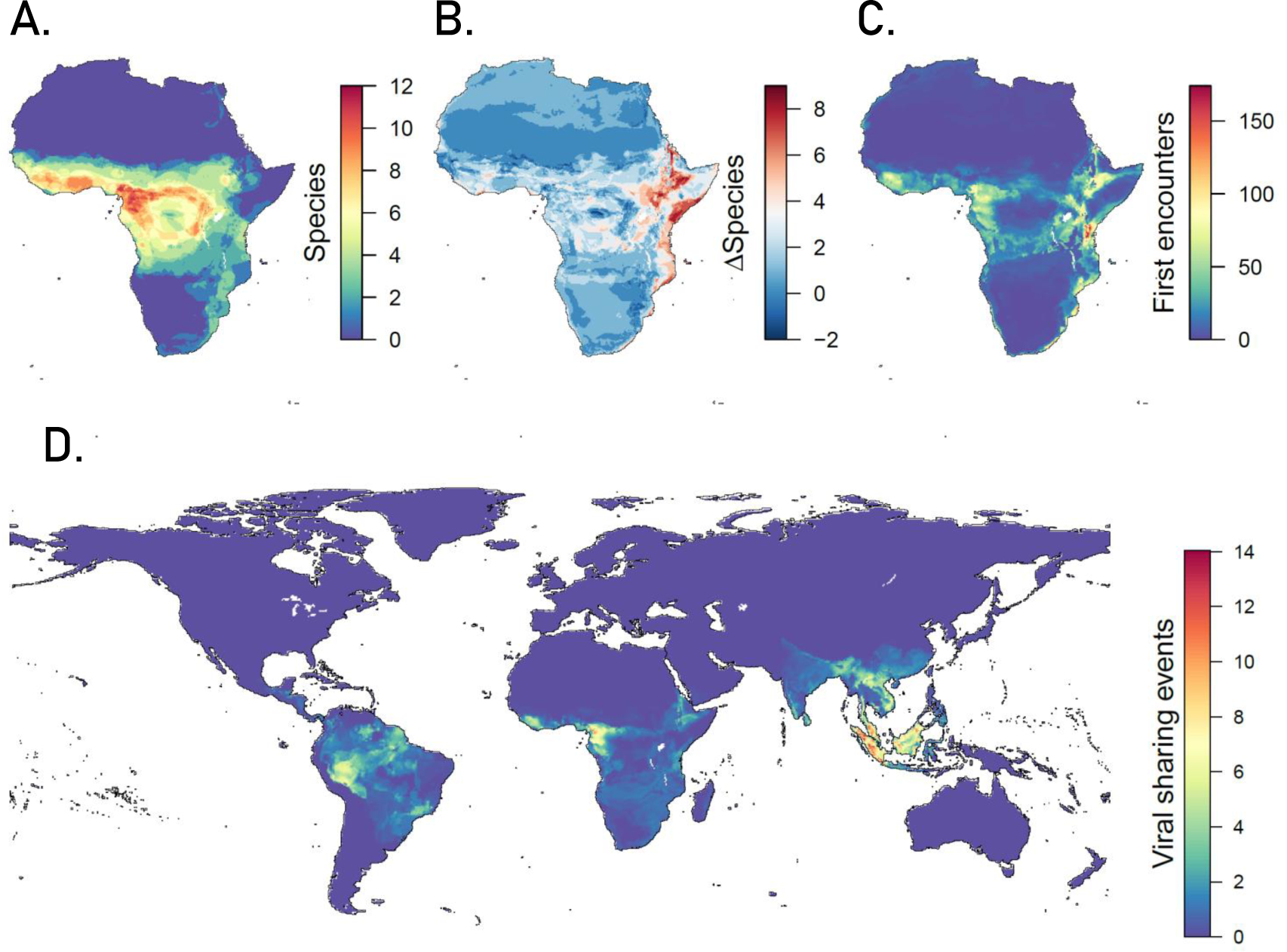
Range expansions will expose naive hosts to zoonotic reservoirs. (A) The predicted distribution of known African hosts of Zaire ebolavirus. (B) The change in richness of these hosts as a result of range shifts (SSP1-RCP 2.6). (C) Projected first encounters with non-Ebola hosts. (D) Bat-primate first encounters are projected to occur globally, producing novel sharing events. Results are averaged across nine global climate models.

### Dispersal drives bats’ disproportionate importance

Species’ dispersal capacity is likely to constrain the ability to move to newly suitable locations, and therefore to limit novel viral sharing. We limited the dispersal potential of flightless species based on an established allometric scaling with body size, trophic rank, and generation time ^32^. Dispersal limits caused substantial reductions in predicted potential range expansions across all scenarios, especially for higher warming scenarios, and therefore drove a reduction in first encounters and novel viral sharing. Even in RCP 2.6 (the scenario with the least warming), limiting dispersal reduced the number of first encounters by 61% (*±* 0.3%), and reduced the associated viral sharing events by 70% (*±* 0.1%) to 4,584 (*±* 52) projected viral sharing events.

Because trophic position and body size determine dispersal capacity, carnivores account for a slightly disproportionate number of first encounters, while ungulates and rodents have slightly fewer first encounters than expected at random (ED Figure 7). Spatial patterns also changed dramatically when dispersal constraints were added, with the majority of first encounters and cross-species viral transmission events occurring in southeast Asia (Figure 1B, ED Figures 4, 6). This viral sharing hotspot is driven disproportionately by bats, because their dispersal was left unconstrained within continents; we made this choice given their exclusion from previous research characterizing the dispersal capacity of range-shifting mammals ^32^, genetic evidence that flight allows bats—and their viruses—to often circulate at continental levels ^46, 47^, and data suggesting that bat distributions are already undergoing disproportionately rapid shifts ^48^. Bats account for nearly 90% of first encounters after constraining dispersal in any climate scenario (RCP 2.6: 88% *±* 0.1%; RCP 8.5: 89% *±* 0.5%), and dominate the spatial pattern, with most of their first encounters restricted to southeast Asia (Figure 2).

Bats’ unique capacity for flight could be an important and previously unconsidered link between climate-driven range shifts and future changes in the mammal virome. Even non-migratory bats can regularly travel hundreds of kilometers within a lifetime, far exceeding what small mammals might be able to cover in 50 years; half of all bat population genetic studies have failed to find any evidence for isolation by distance ^49^. This unique dispersal capacity has inevitable epidemiological implications, with recent evidence suggesting that continental panmixia may be common for zoonotic reservoirs, allowing viral circulation at comparable scales ^46, 47, 50^. Several studies have also identified ongoing rapid range expansions in bat species around the world ^48, 51–58^, with little mention in the broader climate change or emerging disease literature. If flight does allow bats to undergo more rapid range shifts than other mammals, we expect they should drive the majority of novel cross-species viral transmission, and likely bring zoonotic viruses into new regions. This could add an important new dimension to ongoing debate about whether bats are unique in their higher viral richness, higher proportion of zoonotic viruses, or immune adaptations compared to other mammals ^3, 59–63^.

### Impacts on zoonotic viruses and human health

The impacts of climate change on mammalian viral sharing patterns are likely to cascade in future emergence of zoonotic viruses. Among the thousands of expected viral sharing events, some of the highest-risk zoonoses or potential zoonoses are likely to find new hosts. This may eventually pose a threat to human health: the same general rules for cross-species transmission explain spillover patterns for emerging zoonoses ^64, 65^, and the viral species that make successful jumps across wildlife species have the highest propensity for zoonotic emergence ^3, 7, 28^. Just as simian immunodeficiency virus making a host jump from monkeys to chimpanzees and gorillas facilitated the origins of HIV ^66^, or SARS-CoV spillover into civets allowed a bat virus to reach humans ^67^, these kinds of wildlife-to-wildlife host jumps may be evolutionary stepping stones for the *∼*10,000 potentially zoonotic viruses that are currently circulating in mammal hosts ^1^.

To illustrate this problem at the scale of a single pathogen’s “sharing network” (the set of all hosts known or suspected to host the virus, and likely to share with those known hosts), we constructed a sub-network of 13 possible hosts of Zaire ebolavirus (ZEBOV) in Africa, and projected possible first encounters involving these species (Figure 3A-C, ED Figure 8). We project these 13 species to encounter 3,695 (*±*49) new mammals in RCP 2.6, with a modest reduction to 2,627 (*±*44) species when accounting for dispersal limitation, and little variation among climate scenarios (RCP 8.5: 3,529 *±* 47 encounters without dispersal limits; 2,455 *±* 88 with dispersal limits). Even with dispersal limits, these first encounters are predicted to produce almost one hundred new viral sharing events (RCP 2.6: 96 *±* 2; RCP 8.5: 86 *±* 4) that might include ZEBOV, and which cover a much broader part of Africa than the current zoonotic niche of Ebola ^68^. Human spillover risk aside, this could expose several new wildlife species to a deadly virus historically responsible for sizable primate die-offs ^69^. Moreover, for zoonoses like Zaire ebolavirus without known reservoirs, future host jumps—and therefore, the emergence of a larger pool of potential reservoirs covering a greater geographic area (e.g., potential introduction of Zaire ebolavirus to east African mammals)—would only complicate ongoing efforts to trace the sources of spillover and anticipate future emergence ^70, 71^. Ebola is far from unique: with 8,429 *±* 228 first encounters in RCP 2.6 between bats and primates, leading to an expected 110 *±* 4 new viral sharing events even with dispersal limits (Figure 3D; RCP 8.5: 7,326 *±* 667 first encounters, 90 *±* 8 sharing events), many potential zoonoses are likely to experience new evolutionary opportunities because of climate change. Future hotspots of novel mammal assemblages and viral evolution are projected to coincide with areas of high human population density, further increasing vulnerability to potential zoonoses. Potential first encounters are disproportionately likely to occur in areas that are projected to be either human settled or used as cropland and less likely to occur in forests (Figure 2E), despite current literature suggesting that forests harbor most emerging and undiscovered viruses (Figure 4) ^27^. This finding is consistent for bats and non-bats, and may be an accident of geography, but more likely represents the tendency of human settlements to aggregate on continental edges and around biodiversity hotspots ^72^. Regardless of mechanism, we predict that tropical hotspots of novel viral sharing will broadly coincide with high population density areas in 2070, especially in the Sahel, the Ethiopian highlands and the Rift Valley, India, eastern China, Indonesia, and the Philippines (Figure 4). Some European population centers also land in these hotspots; recent emergences in this region like Usutu virus ^73^ highlight that these populations can still be vulnerable, despite greater surveillance and healthcare access. If range-shifting mammals create ecological release for undiscovered zoonoses, populations in any of these areas are likely to be the most vulnerable.

**Figure 4:**
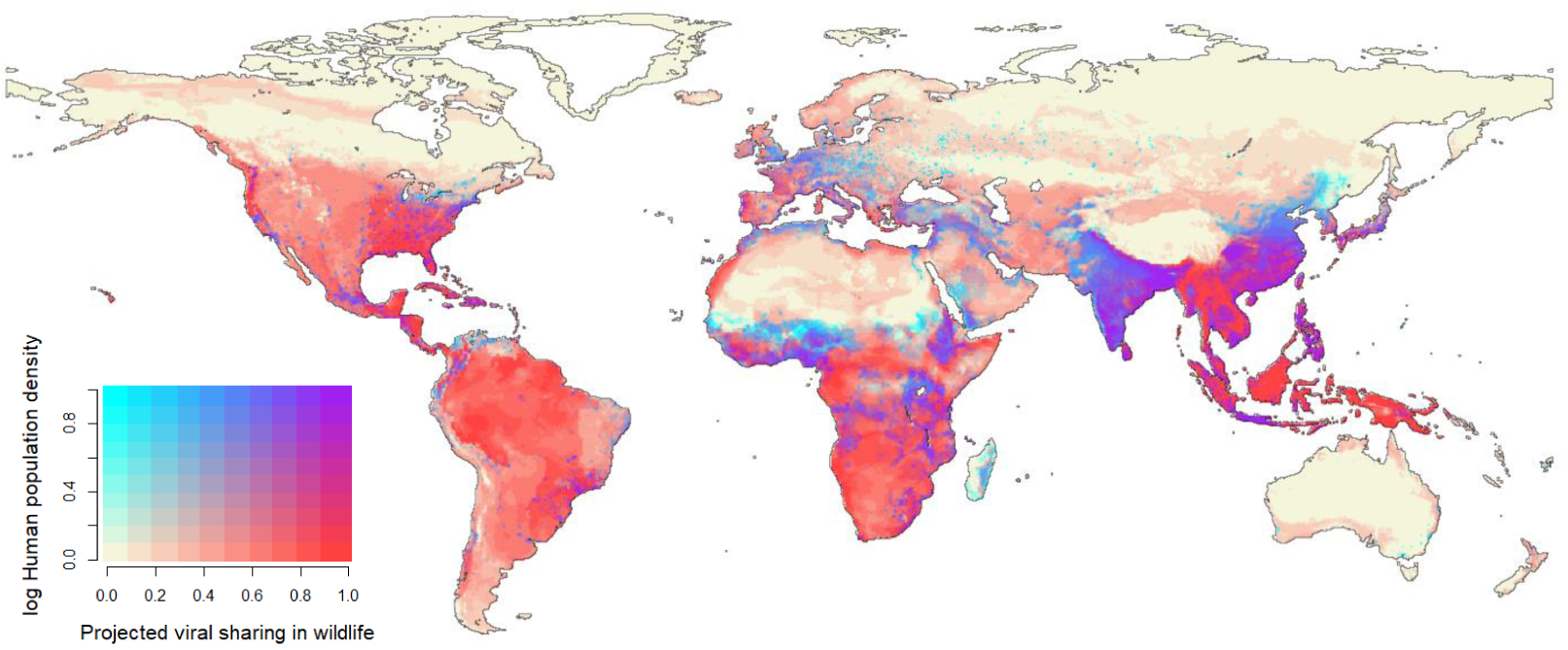
Novel viral sharing events coincide with human population centers. In 2070 (SSP1-RCP 2.6; climate only), human population centers in equatorial Africa, south China, India, and southeast Asia will overlap with projected hotspots of cross-species viral transmission in wildlife. (Both variables are linearly rescaled to 0 to 1.) Results are averaged across nine global climate models.

### Climate change mitigation is insufficient to prevent undesirable outcomes

Whereas most studies agree that climate change mitigation through reducing greenhouse gas emissions will prevent extinctions and minimize harmful ecosystem impacts ^41, 74–77^, our results suggest that mitigation alone cannot reduce the likelihood of climate-driven viral sharing. Instead, the mildest scenarios for global warming appear likely to produce at least as much or even more cross-species viral transmission: when warming is slower, species can successfully track shifting climate optima, leading to more potential for range expansion, and more first encounters. Accounting for dispersal limits, species are projected to experience a median potential loss of 0.3% (*±* 2.5%) of their range in RCP 2.6, with 49.8% (*±* 3.8%) experiencing a net potential increase in range; in contrast, species were predicted to experience a 26.2% (*±* 13.2%) median potential loss in RCP 8.5, and only 30.8% (*±*5.45%) potentially gained any range (ED Figure 3A). In fact, in RCP 8.5, we projected that 261 (*±* 76) species could lose their entire range, with 162 (*±* 53) attributable to dispersal limits alone. As a result, there were 5.4% (*±*1.7%) fewer potential first encounters in RCP 8.5 compared to RCP 2.6, and unexpectedly, a 1.9% (*±* 0.3%) predicted reduction in the connectivity of the future global viral sharing network (ED Figure 3B,D). Overall, our results indicate that a mild perturbation of the climate system could create thousands of new opportunities for viruses to find new hosts. Finally, in a supplemental analysis comparing the present climate to the near past (see Methods and ED Figure 9), we found that if species are already tracking shifting habitats, present-day Africa and the Amazon might already be hotspots of novel cross-species viral transmission, given the warming that has taken place over the last 25 years (*∼* +1*^◦^* C).

We caution that this set of results should not be interpreted as a justification for inaction, or as a possible upside to unmitigated warming, which will be accompanied by mass defaunation, devastating disease emergence, and unprecedented levels of human displacement and global instability ^41, 74–79^. Rather, our results highlight the urgency of better wildlife disease surveillance systems and public health infrastructure as a form of climate change adaptation, even if mitigation efforts are successful and global warming stays below +2°C above pre-industrial levels.

## Conclusions

Our study establishes a macroecological link between climate change and cross-species viral transmission. The patterns we describe are likely further complicated by uncertainties in the species distribution modeling process, including local adaptation or plasticity in response to changing climates, or lack of landscape connectivity preventing dispersal. The projections we make are also likely to be complicated by several ecological factors, including the temperature sensitivity of viral host jumps ^80^; potential independence of vector or non-mammal reservoir range shifts; the possibility that defaunation especially at low elevations might interact with disease prevalence through biodiversity dilution and amplification effects not captured by our models ^81^; or temporal heterogeneity in exposure (hosts might exchange viruses in passing but not overlap by 2070, especially in warmer scenarios). Future work can expand the scope of our findings to other host-parasite systems; our approach, which combines viral sharing models with species distribution modeling approaches for thousands of species, is readily applied to other datasets. Birds have the best documented virome after mammals, and account for the majority of non-mammalian reservoirs of zoonotic viruses ^63^; changing bird migration patterns in a warming world may be especially important targets for prediction. Similarly, with amphibians facing disproportionately high extinction rates due to a global fungal panzootic, and emerging threats like ranavirus causing conservation concern, pathogen exchange among amphibians may be especially important for conservation practitioners to understand ^82^. Finally, marine mammals are an important target given their exclusion here, especially after a recent study implicating reduced Arctic sea ice in novel viral transmission between pinnipeds and sea otters—a result that may be the first proof of concept for our proposed climate-disease link ^83^.

Our study provides the first template for how surveillance could target *future* hotspots of viral emergence in wildlife. In the next decade alone, it may cost at least a billion dollars to comprehensively identify and counteract zoonotic threats before they spread from wildlife reservoirs into human populations ^2^. These efforts are being undertaken during the greatest period of global ecological change recorded in human history, and in a practical sense, the rapid movement of species (and their virome) poses an unexpected challenge for virological research. While several studies have addressed how range shifts in zoonotic reservoirs might expose humans to novel viruses, few have considered the fact that most new exposures will be among wildlife species. The relevance of this process is reinforced by the COVID-19 pandemic, which began only weeks after the completion of this study; the progenitor of SARS-CoV-2 likely originated in southeast Asian horseshoe bats (*Rhinolophus* sp.), and may have spread to humans through an as-yet-unknown bridge host ^84–86^. While we caution against overinterpreting our results as explanatory of the current pandemic or indicative of future pandemic risk—which is largely the product of global health governance, capacity, and preparedness—we note that the global reassortment of mammalian viruses will undoubtedly have a downstream impact on human health (though attribution to climate change will be difficult in any individual case). Tracking spillover into humans is paramount, but so is monitoring of viral transmission in wildlife. Targeting surveillance in future hotspots of cross-species transmission like southeast Asia, and developing norms of open data sharing for the global scientific community, will help researchers identify host jumps early on, ultimately improving our ability to respond to potential threats.

## Methods

In this study, we develop global maps for terrestrial mammals characterizing their habitat use and their ecological niche as a function of climate. We project these into paired climate-land use futures for 2070, with dispersal limitations set by biological constraints for each species. For a final subset of 3,139 species, we predict the probability of viral sharing among species pairs using a model of the mammalian viral sharing network that is trained on phylogenetic relatedness and current geographic range overlaps. With that model, we map the projected hotspots of new viral sharing in different futures. Analysis and visualization code is available on a Github repository (github.com/cjcarlson/iceberg).

### Data

#### Mammal virus data

Our understanding of viral sharing patterns is based on a dataset previously published by Olival *et al.* ^87^. The dataset describes 2,805 known associations between 754 species of mammalian host and 586 species of virus, scraped from the taxonomic data stored in the International Committee on Taxonomy of Viruses (ICTV) database. These data have previously been used in several studies modeling global viral diversity in wildlife ^1, 84, 88^, including a previous study that developed the model of viral sharing we use here ^18^. As that model is reproduced exactly in our study, we have made no further modifications to the data, and more detailed information on data management (e.g., the exclusion of *Homo sapiens* from that analysis) can be found in the Albery *et al.* publication ^18^.

#### Biodiversity data

We downloaded Global Biodiversity Informatics Facility (GBIF: gbif.org) occurrence records for all mammals based on taxonomic names resolved by the IUCN Red List. We developed species distribution models for all 3,870 species with at least three unique terrestrial presence records at a 0.25 degree spatial resolution (approximately 25km by 25 km at the equator). In order to focus on species occurrence, we retained one unique point per 0.25 degree grid cell. This spatial resolution was chosen to match the available resolution of land use change projections (see below). Spatial and environmental outliers were removed based on Grubb outlier tests ^89^. To implement the Grubb outlier tests for a given species we defined a distance matrix between each record and the centroid of all records (in both environmental or geographic space, respectively) and determined whether the record with the largest distance was an outlier with respect to all other distances, at a given statistical significance (*p* = 1*e −* 3, in order to exclude only extreme outliers). If an outlier was detected it was removed and the test was repeated until no additional outliers were detected.

#### Climate and land use data

Climate and land use data were compiled from WorldClim 2 ^90^ and the Land Use Harmonization 2 (LUH2) project ^91^ respectively, for both baseline conditions (operationalized as 1970-2000 for the climate data, 2015 for land use, and 2020 for dispersal limits; see “The effect of recent warming” for an interrogation of the difference between climate baselines and actual present-day climate) and a half-century in the future (operationalized as 2061-2080 for climate, 2070 for land use, and 2070 for dispersal).

The WorldClim dataset is widely used in ecology, biodiversity, and agricultural projections of potential climate change impacts. WorldClim makes data available for current and future climates in the form of 19 pre-processed bioclimatic variables (Bioclim: BIO1-19). In order to reduce collinearity among climate variables in the species distribution models, we selected five Bioclim variables from the full set of 19 Bioclim variables: mean annual temperature (BIO1), temperature seasonality (BIO4), annual precipitation (BIO12), precipitation seasonality (coefficient of variation; BIO15), and precipitation of the driest quarter (BIO17). This is the largest set of Bioclim variables possible that keeps their correlation over a global extent suitably low (r *<* 0.7). The Bioclim variables for the historical climate are the mean from 1970-2000, and those for the future climate are the mean from 2060-2080.

To account for model uncertainty in climate projections, we used projections for future climates from all nine global climate models (GCMs) currently available on WorldClim 2 and participating in the Coupled Model Intercomparison Project 6 (CMIP6), the most recent generation of climate models: BCC-CSM2-MR, CNRM-CM6-1, CNRM-ESM2-1, CanESM5, GFDL-ESM4, IPSL-CM6A-LR, MIROC-ES2L, MIROC6, and MRI-ESM2-0. These nine GCMs encompass a wide range of effective climate sensitivities from 2.6K (MIROC6) to 5.6K (CanESM5) compared with a range of 1.8-5.6K across 27 CMIP6 models and 2.1-4.7K for CMIP5 ^92^. Temperature and precipitation for future climates have been downscaled and bias-corrected by WorldClim 2 using a change factor approach. The multi-year average of the GCM output for minimum temperature, maximum temperature and total precipitation is calculated for each month of the simulated historical and future period, and the absolute (for temperature) or proportional (for precipitation) difference in these values is then calculated, resulting in climate anomalies which are then applied to the 10-minute spatial resolution observed historical dataset ^90, 93^. WorldClim 2 then calculates Bioclim variables based on these downscaled and bias-corrected data. This approach makes the assumption that the change in climate is relatively stable across space (that is, has high spatial autocorrelation). We downloaded the five pre-processed Bioclim variables for all nine GCMs at 10 minutes spatial resolution from WorldClim 2 ^90^, and aggregated with bilinear interpolation to 0.25 degree spatial resolution (approximately 25km at the equator) to match with the LUH2 land use data resolution.

Historical land-use data for 2015 and projected land-use data for 2070 were obtained from the Land Use Harmonization 2 (LUH2) project at 0.25 degree spatial resolution ^91, 94^. The LUH2 data reconstructs and projects changes in land use among twelve categories: primary forest, non-forested primary land, potentially forested secondary land, potentially non-forested secondary land, managed pasture, rangeland, cropland (four types), and urban land. To capture species’ habitat preferences, we downloaded data for all 3,870 mammal species from the IUCN Habitat Classification Scheme (version 3.1) and mapped the 104 unique IUCN habitat classifications onto the twelve land use types present in the LUH2 dataset following Powers *et al.* ^95^ (Supplementary Table 1).

Finally, we downloaded global population projections from the SEDAC Global 1-km Down-scaled Population Base Year and Projection Grids Based on the SSPs version 1.0 ^96^, and selected the year 2070 for RCP 2.6 (see “Climate and land use futures”). These data are downscaled to 1km from a previous dataset at 7.5 arcminute resolution ^97^. We aggregated 1 km grids up to 0.25 degree grids for compatibility with other layers, again using bilinear interpolation.

#### Additional data

A handful of smaller datasets were incidentally used throughout the study. These included the IUCN Red List, which was used to obtain species taxonomy, range maps, and habitat preferences ^98^; the US Geological Survey Global Multi-resolution Terrain Elevation Data 2010 dataset, which was used to derive a gridded elevation in meters at *∼*25km resolution; and a literature-derived list of suspected hosts of Ebola virus ^30^.

### Mapping species distributions

We developed species distribution models for a total of 3,870 species in this study, divided into two modeling pipelines based on data availability (ED Figures 10, 11).

#### Poisson point process models

For 3,088 species with at least 10 unique presence records, Poisson point process models (PPMs), a method closely related to maximum entropy species distribution models (MaxEnt), were fit using regularized downweighted Poisson regression ^99^ with 20,000 background points, using the R package glmnet ^100, 101^. The spatial domain of predictions was chosen based on the continent(s) where a species occurred in their IUCN range map; as a final error check, species ranges were constrained to a 1,000 km buffer around their IUCN ranges. We trained species distribution models on current climate data using the WorldClim 2 data set ^90^, using the five previously-specified Bioclim variables.

To reduce the possibility of overfitting patterns due to spatial aggregation, we used spatially stratified cross validation. Folds were assigned by clustering records based on their coordinates and splitting the resulting dendrogram into 25 groups. These groups were then randomly assigned to five folds. (If species had fewer than 25 records, a smaller number of groups was used based on sample size, and these were split into five folds.) This flexible approach accounts for variation in the spatial scale of of aggregation among species by using the cluster analysis. By splitting into 25 groups initially (rather than 5) we obtain better environmental coverage (at least on average) within a fold and minimize the need to extrapolate for withheld predictions.

Linear (all species), quadratic (species with >100 records), and product (species with >200 records) features were used. Positive coefficients of quadratic features are not allowed (i.e. all have an upper bound of 0 in the model-fitting process), to avoid the undesirable effect of increasing suitability predictions at range edges. The regularization parameter was determined based on 5-fold cross-validation with each fold, choosing a value 1 standard deviation below the minimum deviance ^102^. This resulted in five models per species which were then combined in an unweighted ensemble. Continuous predictions of the ensemble were converted to binary presence/absence predictions by choosing a threshold based on the 5th percentile of the ensemble predictions at training presence locations.

When models were projected into the future, we limited extrapolation to 1 standard deviation beyond the data range of presence locations for each predictor. This decision balances a small amount of extrapolation based on patterns in a species niche with limiting the influence of monotonically increasing marginal responses, which can lead to statistically unsupported (and likely biologically unrealistic) responses to climate.

#### Range bagging models

For an additional 783 rare species (3 to 9 unique points on the 25 km grid), we produced species distribution models with a simpler range bagging algorithm, a stochastic hull-based method that can estimate climate niches from an ensemble of underfit models ^103, 104^, and is therefore well suited for smaller datasets. From the full collection of presence observations and environmental variables range-bagging proceeds by randomly sampling a subset of presences (proportion *p*) and a subset of environmental variables (*d*). From these, a convex hull around the subset of points is generated in environmental space. The hull is then projected onto the landscape with a location considered part of the species range if its environmental conditions fall within the estimate hull. The subsampling is replicated *N* times, generating *N* ‘votes’ for each cell on the landscape. One can then choose a threshold for the number of votes required to consider the cell as part of the species’ range to generate the binary map used in our downstream analyses. Based on general guidelines in ^103^ we chose *p* = 0.33, *d* = 2, and *N* = 100. We then chose the voting threshold to be 0.165 (=0.33/2) because this implies that the cell is part of the range at least half the time for each subsample. Upon visual inspection, this generally lead to predictions that were very conservative about inferring that unsampled locations were part of a species distribution. The same environmental predictors and ecoregion-based domain selection rules were used for range bagging models as were used for the point process models discussed above. This hull-based approach is particularly valuable for poorly sampled species which may suffer from sampling bias because bias within niche limits has little effect on range estimates.

#### Model validation and limitations

PPM models performed well, with a mean test AUC under 5 fold cross-validation (using spatial clustering to reduce inflation) of 0.78 (s.d. 0.14). The mean partial AUC evaluated over a range of sensitivity relevant for SDM (0.8-0.95) was 0.81 (s.d. 0.09). The mean sensitivity of binary maps used to assess range overlap (based on the 5% training threshold used to make a binary map) was 0.90 (s.d. 0.08). Range bagging models were difficult to meaningfully evaluate because they were based on extremely small sample sizes (3-9). The mean training AUC (we did not perform cross-validation due to small sample size) was 0.96 (s.d. 0.09). The binary maps had perfect sensitivity (1) because the threshold used to make them was chosen sufficiently low to include the handful of known presences for each species. One way to assess how well we inferred the range for these species is to quantify how much of the range was estimated based on our models, based on the number of (10km) cells predicted to be part of the species range even when it was not observed there. The mean number of cells inferred to contain a presence was 254 (s.d. 503); however, the distribution is highly right skewed with a median of 90. This indicates that the range bagging models were typically relatively conservative about inferring ranges for poorly sampled species.

Although our models performed well, we note that researchers should approach the interpretation of species distribution models (SDMs) with a certain degree of caution. Even adhering to best practices, many SDM methods are sensitive to subjective user-end choices that influence model performance, transferrability, and interpretability. Some of those choices may have marginally affected the patterns we document in this study. For example, to quantify our results’ resilience to the choice of threshold, we constructed pairwise overlaps for the current rasters of all species across three habitat suitability thresholds (1%, 5%, and 10%). We did this using the climate projections, the IUCN-clipped climate projections, and the land use-and IUCN-clipped projections (see below sections), such that there were nine total replicates, only one of which (IUCN-and land use-clipped 5% threshold) was used in our main analyses. We fitted the proportional overlap between each species pair across all nine replicates in a linear mixed model with the identity of the species pair and the thresholding replicate as random effects, to quantify the variance associated with the choice of processing pipeline compared to the variance associated with the species pair itself. We also examined the mean proportional overlap across the nine replicates. Our linear mixed model examining the variance associated with thresholding pipeline found that thresholding accounted for only 2.2% of the variance in proportional overlap, in contrast to the 72.3% accounted for by the identity of the species pair. Furthermore, there was very little difference observed in the mean proportional overlap and the number of overlapping species across thresholds. These results demonstrate that the choice of thresholding had an impact on the results of our analysis, but an extremely marginal one, and we expect similar results would be found for other choices like variable set reduction, model calibration, the resolution of predictor data, and the processing of point occurrence data.

Finally, we note that while many factors besides climate are ignored by our models, such as biotic interactions or animal social behavior, our models are tailored to our aim: predicting hotspots of elevated risk under climate change. In our application, correctly predicting presences is more important than incorrect prediction of absences, because we are focused on the potential for novel species overlap. We cannot say whether that overlap *will* happen, based on the multiple factors besides climate that influence distributions and range shifts, but we can say with confidence - based on robust current niche estimates, validated with spatially stratified cross-validation, and biologically-grounded estimates of dispersal capacity - where risk would be elevated in accordance with our simulations.

#### Habitat range and land use

To capture species’ habitat preference, we collated data for all 3,870 mammal species from the IUCN Habitat Classification Scheme (version 3.1). We then mapped 104 unique IUCN habitat classifications onto the twelve land use types present in the LUH2 dataset. For 962 species, no habitat data was available, or no correspondence existed between a land type in the IUCN scheme and our land use data; for these species, land use filters were not used. Filtering based on habitat was done as permissively as possible: species were allowed in current and potential future ranges to exist in a pixel if any non-zero percent was assigned a suitable habitat type; almost all pixels contain multiple habitats. In some scenarios, human settlements cover at least some of a pixel for most of the world, allowing synanthropic species to persist throughout most of their climatically-suitable range. For those with habitat data, the average reduction in range from habitat filtering was 7.6% of pixels.

### Predicting future species distributions

We modeled a total of 136 future scenarios, produced by the four paired climate-land use change pathways replicated across nine global climate models (with one, GFDL-ESM4, only available for two climate scenarios: RCP 2.6 and RCP 7.0; see below), modified by two optional filters on species ranges (habitat preferences and dispersal limits). The full matrix of possible scenarios captures a combination of scenario uncertainty about global change and epistemological uncertainty about how best to predict species’ range shifts. By filtering potential future distributions based on climate, land use, and dispersal constraints, we aimed to maximize realism; our predictions were congruent with extensive prior literature on climate- and land use-driven range loss ^95, 105, 106^.

#### Climate and land use futures

We considered four possible scenarios for the year 2070 each based on a pairing of the Representative Concentration Pathways (RCPs) and the Shared Socioeconomic Pathways (SSPs). RCP numbers (e.g., 2.6 or 4.5) represent Watts per square meter of additional radiative forcing by the end of the century, while SSPs describe alternate possible pathways of socioeconomic development and demographic change. As pairs, SSP-RCP scenarios describe alternative futures for global socioeconomic and environmental change. Not all SSP-RCP scenario combinations in the “scenario matrix” are realistically possible ^107^. For example, in the vast majority of integrative assessment models, decarbonization cannot be achieved fast enough in the SSP5 scenario to achieve RCP 2.6.

We used four SSP-RCP combinations: SSP1-RCP2.6, SSP2-RCP4.5, SSP3-RCP7.0, and SSP5-RCP8.5. We selected these four scenarios because they span a wide range of plausible global change futures, and serve as the basis for climate model projections in the Scenario Model Intercomparison Project for the newest generation of global climate models (CMIP6) ^31^. SSP1-RCP2.6 is a scenario with low population growth, strong greenhouse gas mitigation and land use change (especially an increase in global forest cover), which makes global warming likely less than 2*^◦^* C above pre-industrial levels by 2100; SSP2-RCP4.5 has moderate land use change and greenhouse gas mitigation with global warming of around 2.5*^◦^* C by 2100; SSP3-RCP7.0 has high population growth, substantial land use change (especially a decrease in global forest cover) and very weak greenhouse gas mitigation efforts with global warming of around 4*^◦^* C by 2100; and SSP5-RCP8.5 is the highest warming scenario with less decrease in forest cover than SSP3 but more substantial increases in coal and other fossil fuel usage leading to more than 4*^◦^* Cwarming by 2100 ^31, 108–110^.

#### Climate model uncertainty

To identify the contribution of climate model uncertainty and its propagation through our analysis, we used all nine selected GCMs from CMIP6 and produced multi-model averages for all main text figures. For all of the main text statistics, we present each multi-model mean with a standard deviation across the nine global climate models. We also compared the first encounters from the two models with the highest (CanESM5) and lowest (MIROC6) effective climate sensitivity in the available CMIP6 set on WorldClim (ED Figure 5) ^92^. We also present the map of first encounters and novel viral sharing in each GCM run for each RCP, accounting for both climate and land use change, with the full dispersal and limited dispersal scenario, in Supplementary Figures 1-18.

#### Limiting dispersal capacity

Not all species can disperse to all environments, and not all species have equal dispersal capacity—in ways likely to covary with viral sharing properties. We follow a rule proposed by Schloss *et al.* ^32^, who described an approximate formula for mammal range shift capacity based on body mass and trophic position. For carnivores, the maximum distance traveled in a generation is given as *D* = 40.7*M*^0.81^, where *D* is distance in kilometers and *M* is body mass in kilograms. For herbivores and omnivores, the maximum is estimated as *D* = 3.31*M*^0.65^.

We used mammalian diet data from the EltonTraits database ^111^, and used the same cutoff as Schloss to identify carnivores as any species with 10% or less plants in their diet. We used body mass data from EltonTraits in the Schloss formula to estimate maximum generational dispersal, and converted estimates to annual maximum dispersal rates by dividing by generation length, as previously estimated by another comprehensive mammal dataset ^112^. We multiply by 50 years (from 2020 as the present to 2070) and use the resulting distance as a buffer around the original range map, and constrain possible range shifts within that buffer. For 420 species with missing data in one of the required sources, we interpolated dispersal distance based on the closest relative in our supertree with a dispersal velocity estimate.

Qualified by the downsides of assuming full dispersal ^113^, we excluded bats from the assumed scaling of dispersal limitations. The original study by Schloss *et al.* ^32^ chose to omit bats entirely, and subsequent work has not proposed any alternative formula. Moreover, the Schloss formula performs notably poorly for bats: for example, it would assign the largest bat in our study, the Indian flying fox (*Pteropus giganteus*), a dispersal capacity lower than that of the gray dwarf hamster (*Cricetulus migratorius*). Bats were instead given full dispersal in all scenarios: given significant evidence that some bat species regularly cover continental distances ^46, 47^, and that isolation by distance is uncommon within many bats’ ranges ^49^, we felt this was a defensible assumption for modeling purposes. Moving forward, the rapid range shifts already observed in many bat species (see main text) could provide an empirical reference point to fit a new allometric scaling curve (after standardizing those results for the studies’ many different methodologies). A different set of functional traits likely govern the scaling of bat dispersal, chiefly the aspect ratio (length:width) of wings, which is a strong predictor of population genetic differentiation ^49^. Migratory status would also be important to include as a predictor although here, we exclude information on long-distance migration for all species (due to a lack of any real framework for adding that information to species distribution models in the literature).

### Explaining spatial patterns

To explore the geography of novel assemblages, we used linear models that predicted the number of first encounters (novel overlap of species pairs) at the 25km level (*N* = 258, 539 grid cells). Explanatory variables included: richness (number of species inhabiting the grid cell in our predicted current ranges for the given scenario); elevation in meters (derived from the US Geological Survey Global Multi-resolution Terrain Elevation Data 2010 dataset); and the pre-dominant land cover type for the grid cell. We simplified the classification scheme for land use types into five categories for these models (human settlement, cropland, rangeland and pasture, forest, and unforested wildland), and assigned pixels a single land use type based on the maximum probability from the land use scenarios. We fit a model for each scenario and pair of biological assumptions; because of the large effect bats had on the overall pattern, we retrained these models on subsets of encounters with and without a bat species involved. To help model fitting, we log(x+1)-transformed the response variable (number of overlaps in the pixel) and both continuous explanatory variables (meters of elevation above the lowest point and species richness). Because some elevation values were lower than 0 (i.e., below sea level), we treated elevation as meters above the lowest terrestrial point rather than meters above sea level to allow us to log-transform the data.

### Viral sharing models

#### Criteria for species’ inclusion

Of the 3,870 species for which we generated distribution models, 103 were aquatic mammals (cetaceans, sirenians, pinnipeds, and sea otters), and 382 were not present in the mammalian supertree that we used for phylogenetic data ^114^. These species, and the associated species distribution models, were excluded from the analysis. Aquatic species were removed using a two-filter approach, by first cross-referencing with Pantheria ^115^, and second by checking no species only had non-aquatic habitat use types (see “Habitat range and land use”). We also excluded 246 monotremes and marsupials because the shape of the supertree prevented us from fitting satisfactory GAMM smooths to the phylogeny effect, leaving 3,139 non-marine placental mammals with associated phylogenetic data.

#### Generalized additive mixed models

We used a previously-published model of the phylogeography of viral sharing patterns to make predictions of future viral sharing ^18^. This model was based on an analysis of 510 viruses shared between 682 mammal species ^3^, and predicted the probability that a pair of mammal species will share a virus given their geographic range overlap and phylogenetic relatedness. The original study uncovered strong, nonlinear effects of spatial overlap and phylogenetic similarity in determining viral sharing probability, and simulating the unobserved global network using these effect estimates capitulated multiple macroecological patterns of viral sharing.

In the original study, a Generalized Additive Mixed Model (GAMM) was used to predict virus sharing as a binary variable, based on (1) geographic range overlap; (2) phylogenetic similarity; and (3) species identity as a multi-membership random effect. The phylogeographic explanatory variables were obtained from two broadly available, low-resolution data sources: pairwise phylogenetic similarity was derived from a mammalian supertree previously modified for host-pathogen studies ^3, 114^, with similarity defined as the inverse of the cumulative branch length between two species, scaled to between 0 and 1. Geographic overlap was defined as the area of overlap between two species’ IUCN range maps, divided by their cumulative range size ^116^.

We first retrained the GAMMs from ^18^ on the pairwise overlap matrix of species distribution models generated for this study, so that present predictions would be comparable with potential future distributions. Of the 3,139 species in our reduced dataset, 544 had viral records in our viral sharing dataset and shared with at least one other mammal, and were used to retrain the GAMM from ^18^. To check the performance of the GAMM, we predicted sharing patterns with a) only random effects, b) only fixed effects, and c) with both. To extend predictions to the the full set of mammals, we generated random effects for out-of-sample species by drawing from the fitted distribution of species-level effects. (Predicting without these random effects underestimates species variance, resulting in mean sharing of 0.02 rather than the observed 0.06). The mean sharing value across these predictions closely approximated observed sharing probability (*∼* 0.06).

We note that this model uses citation counts to correct for sampling bias, an imperfect method but one that leads to strong validation performance on an independently-compiled dataset of host-virus associations, which carries a different set of biases. However, it is still possible that sampling bias in host-virus datasets like the Olival *et al.* dataset could artificially inflate the signal of phylogeography in viral sharing, if researchers investigating a noteworthy viral detection then preferentially sample closely-related host species in the immediate area. It is unlikely these effects would bias our results in a particular direction, but accounting for these biases should at least involve some acknowledgement that cross-species transmission is challenging to predict. (See the Albery *et al.* study’s Discussion for a more in-depth treatment of sampling bias effects.)

#### Model validation and limits

Compared to the current viral sharing matrix, the model performs well with only fixed effects (AUC = 0.80) and extremely well with both fixed and random effects (AUC = 0.93). The model explained a very similar proportion of the deviance in viral sharing to that in Albery *et al.* ^18^ (44.5% and 44.8%, respectively).

In practice, several unpredictable but confounding factors could affect the reliability of this model as a forecasting tool, including temperature sensitivity of viral evolution in host jumps ^80^, or increased susceptibility of animals with poorer health in lower-quality habitat or unfavorable climates. Moreover, once viruses can produce an infection, their ability to transmit *within* a new species is an evolutionary race between mutation and recombination rates in viral genomes, host innate and adaptive immunity, virulence-related mortality, and legacy constraints of coevolution with prior hosts and vectors ^64, 65^. But data cataloging these precise factors are hardly comprehensive for the hundreds of zoonotic viruses, let alone for the thousands of undescribed viruses in wildlife. Moreover, horizontal transmission is not necessary for spillover potential to be considered significant; for example, viruses like rabies or West Nile virus are not transmitted within human populations but humans are still noteworthy hosts.

#### Mapping opportunities for sharing

We used the GAMM effect estimates to predict viral sharing patterns across the 3,139 mammals with associated geographic range and phylogenetic data, for both the present and future scenarios. By comparing current and future sharing probabilities for each of the four global change scenarios, we estimated which geographic and taxonomic patterns of viral sharing would likely emerge. We separately examined patterns of richness, patterns of sharing probability, and their change (i.e., future sharing probability - current sharing probability, giving the expected probability of a novel sharing event).

A subset of the mammals in our dataset were predicted to encounter each other for the first time during range shifts. For each of these pairwise first encounters, we extracted the area of overlap in every future scenario, and assigned each overlap a probability of sharing from the mean GAMM predictions and mapped the mean and cumulative probability of a new sharing event happening in a given geographic pixel.

#### Case study on Zaire ebolavirus

For a case study in possible significant cross-species transmission, we compiled a list of known hosts of Zaire ebolavirus (ZEBOV), a zoonosis with potentially high host breadth that has been known to cause wildlife die-offs, but has no known definitive reservoir. Hosts were taken from two sources: the training dataset on host-virus associations ^3^, and an additional dataset of filovirus testing in bats ^30^. In the latter case, any bats that have been reported antibody positive or PCR-positive for ZEBOV were included. A total of 19 current “known hosts” were selected. We restricted our analysis to the 13 hosts from Africa, because there is no conclusive evidence that Zaire ebolavirus actively circulates outside Africa; although some bat species outside Africa have tested positive for antibodies to ZEBOV, this is likely due to cross-reactivity with other undiscovered filoviruses ^30, 117, 118^. We used the 13 African hosts to predict possible first encounters in all scenarios (ED Figure 8), and mapped the current richness of ZEBOV hosts, the change in host richness by 2070, and the number of first encounters (Figure 3).

#### Overlap with human populations

To examine the possibility that hotspots of cross-species transmission would overlap with human populations, we used SEDAC’s global population projections version 1.0 for the year 2070 ^96^. We aggregated these to native resolution, for each of the four SSP paired with the native RCP/SSP pairing for the species distribution models. In Figure 4 we present the population projections for SSP1, which pairs with RCP 2.6.

### The effect of recent warming

Like many studies that employ species distribution modeling, our study uses a definition of the “present” that embodies a slight cognitive dissonance with recent warming ^119^. The World-Clim2 dataset captures the mean climate between 1970 and 2000, but the climate at the time of writing has already warmed substantially compared to this baseline. While we employ this loose definition of “present day” throughout, we note that the actual present climate is substantially warmer, and therefore might be expected to already be experiencing the turnover in viral sharing that we describe throughout.

As a final supplementary analysis, we interrogated the effect of recent climate change on the world we live in today, which is already substantially warmer than pre-industrial temperatures. To do so, we repeated the analysis in its entirety – minus steps constraining species ranges with either the IUCN range maps or dispersal limits – using the ERA5 reanalysis product with monthly averaged data ^120^. We trained species distribution models based on a recent climate baseline (1981-1995), and projected their ranges to the present day (2005-2019), using two time slices (1991 and 2015) positioned equally in the climate intervals. We set dispersal limits for species as we did in the main analysis, but for this 25-year period.

Using these data to repeat the analysis, we found that there were a projected total of 52,463 first encounters (with 34,254 including at least one bat species), amounting to a total of 1,043 viral sharing events. First encounters and viral sharing events were located mostly in Africa and the Amazon (ED Figure 9). We caution that these results–particularly the number of encounters and sharing events–should not be interpreted as the same “units” as the main analysis, given that they are calibrated to an entirely different climate reconstruction.

## Acknowledgements

This paper is the culmination of several years of idea development and owes special thanks to many people, including the entire Bansal Lab, Laura Ward Alexander, Kevin Burgio, Eric Dougherty, Romain Garnier, Wayne Getz, Peta Hitchens, Christine Johnson, and Isabel Ott. We especially thank Laura Alexander for sharing bat filovirus testing sources used to compile the Ebola sub-network. Thanks are also extended to José Hidasi-Neto for publicly-available data visualization code. CJC was supported by the Georgetown Environment Initiative and the National Socio-Environmental Synthesis Center (SESYNC) under funding received from the National Science Foundation DBI-1639145. CJC, GFA, and EAE were supported by funding to the Verena Consortium including NSF BII 2021909 and a grant from Institut de Valorisation des Données (IVADO). CM acknowledges funding from National Science Foundation grant DBI-1913673. EAE, KJO, and NR were supported by the United States Agency for International Development (USAID) Emerging Pandemic Threats PREDICT project.

## Author Contributions

CJC and GFA conceived the study. CM, CJC, and CHT developed species distribution models; GFA, EAE, KJO, and NR developed the generalized additive models. GFA, CJC, and CMZ integrated the predictions of species distributions and viral sharing patterns and designed visualizations. All authors contributed to the writing of the manuscript.

---

**Extended Data Figure 1:**
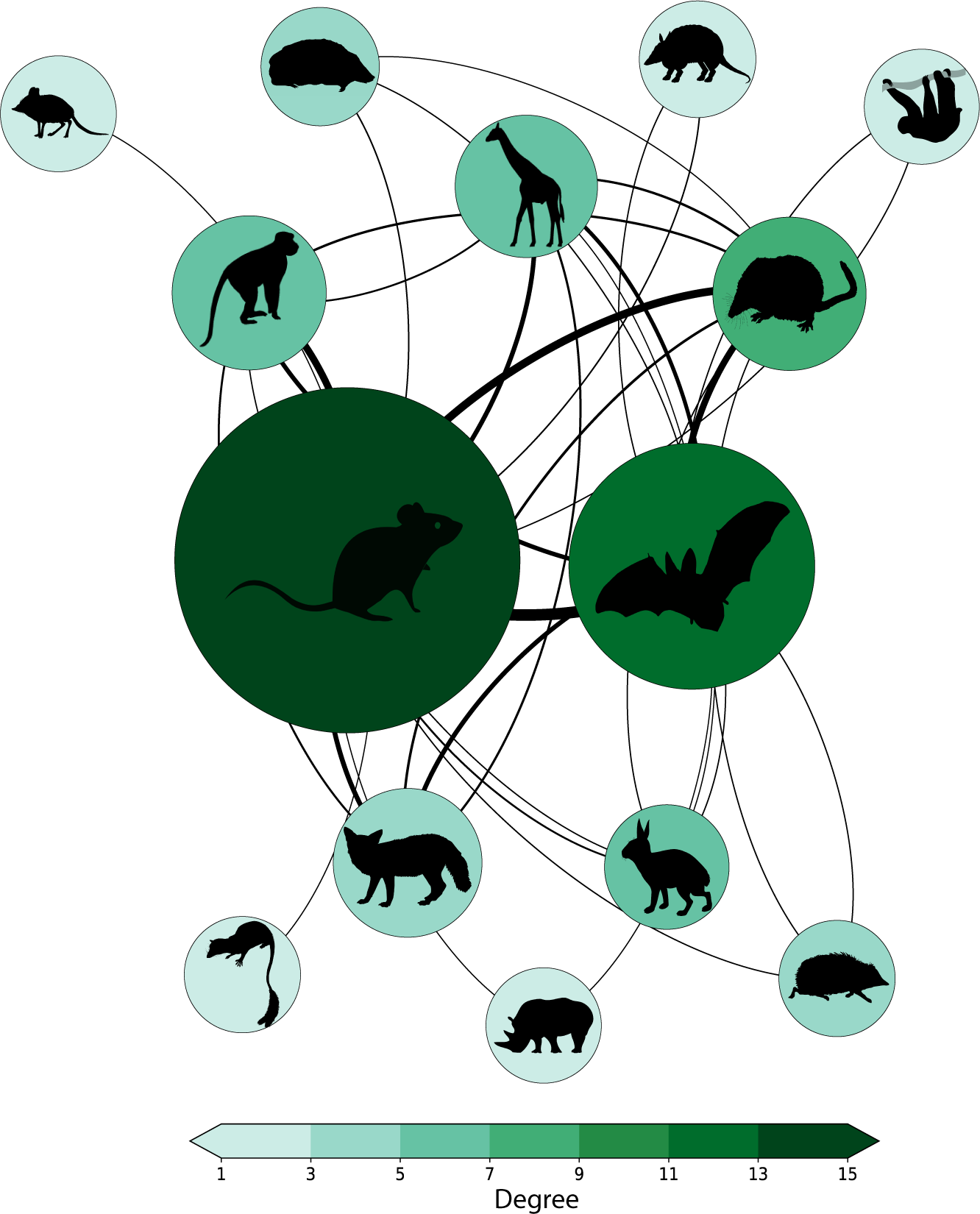
The mammal-virus network. The present-day viral sharing network by mammal order inferred from modeled pairwise predictions of viral sharing probabilities. Edge width denotes the expected number of shared viruses (the sum of pairwise species-species viral sharing probabilities), with most sharing existing among the most speciose and closely-related groups. Edges shown in the network are the top 25% of links. Nodes are sized by total number of species in that order in the host-virus association dataset, color is scaled by degree.

**Extended Data Figure 2:**
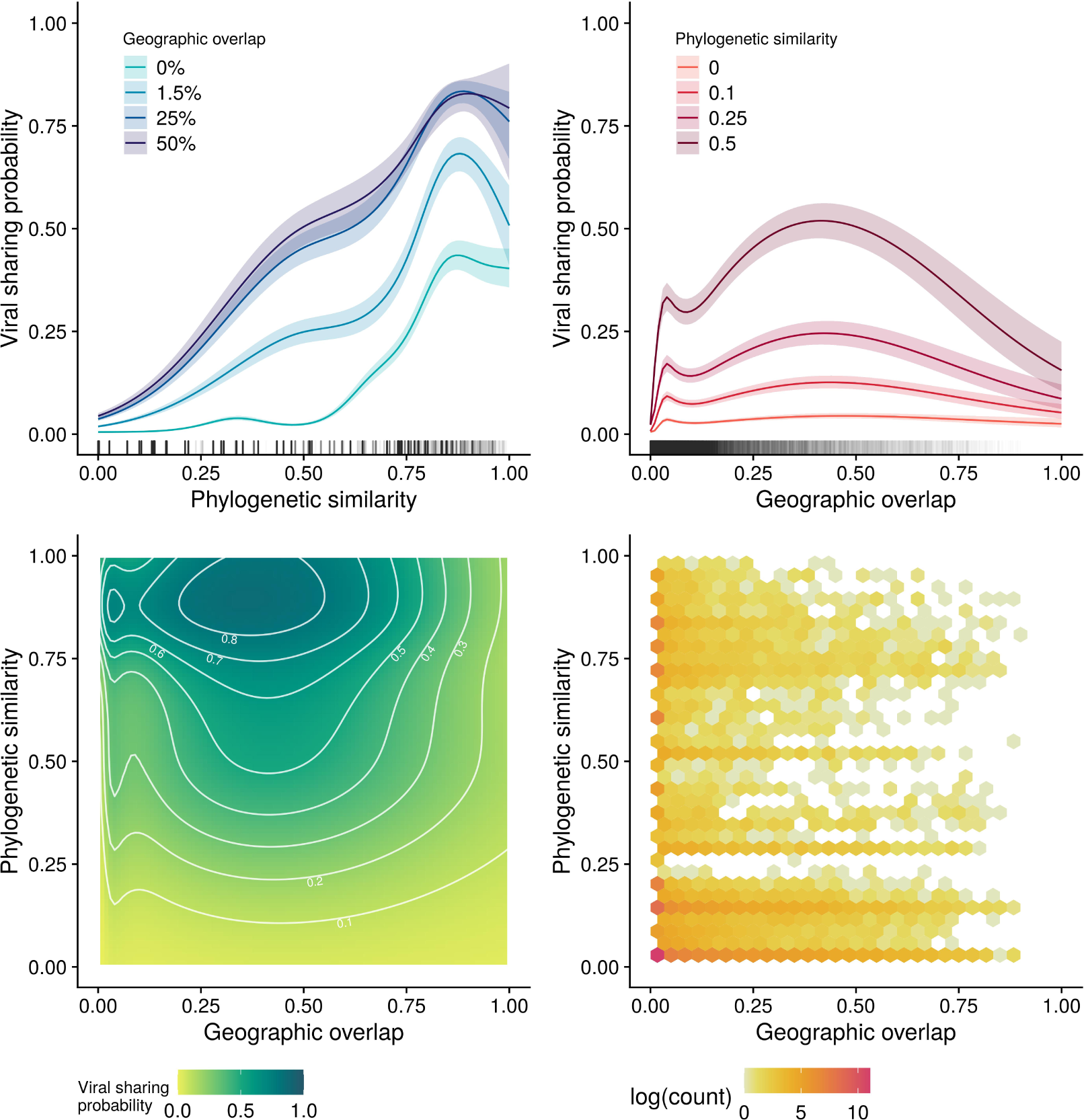
Predicted phylogeographic structure of viral sharing. Phylogeographic prediction of viral sharing using a generalized additive mixed model. Viral sharing increases as a function of phylogenetic similarity (A) and geographic overlap (B), which have strong nonlinear interactions, shown in the contour map of joint effects (C). White contour lines denote 10% increments of sharing probability. Declines at high values of overlap may be an artefact of model structure and low sampling in the upper levels of geographic overlap, shown in a hexagonal bin chart of the raw data distribution (D).

**Extended Data Figure 3:**
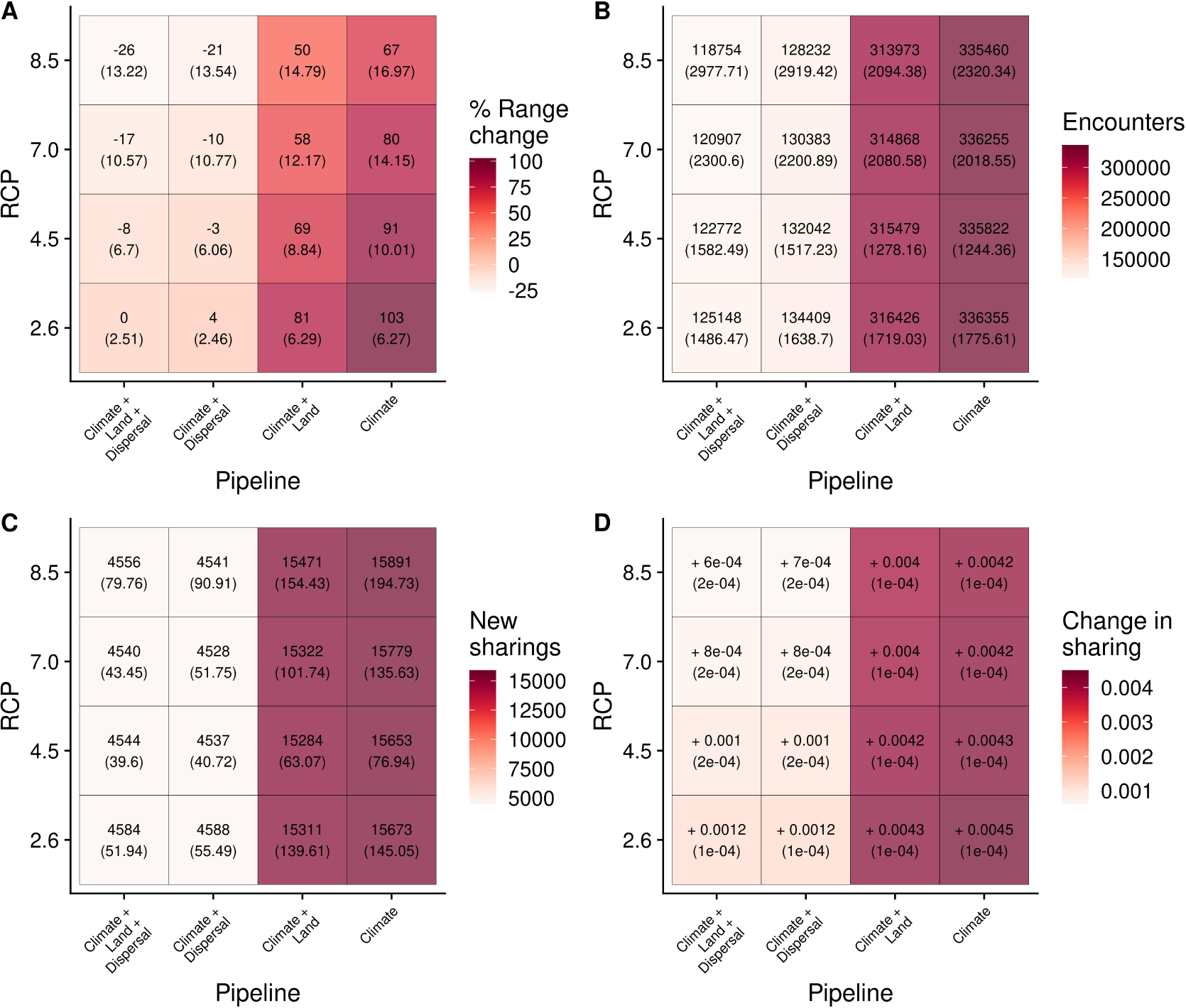
Outcomes by model formulation and climate change scenario. Heatmaps displaying predicted changes across model formulations. (A) Range expansions were highest in non-dispersal-limited scenarios and in scenarios with lower levels of global warming. (B) The number of predicted first encounters was higher in non-dispersal-limited scenarios and in scenarios with lower levels of global warming. (C) The number of expected new viral sharing events was higher in non-dispersal-limited scenarios and in more severe RCPs. (D) The overall change in sharing probability (connectance) across the viral sharing network between the present day and the future scenarios; absolute change is minimal but positive across all scenarios, being greatest in non-dispersal-limited scenarios and in scenarios with lower levels of global warming. Results are averaged across nine global climate models, with standard deviation indicated in parentheses underneath main statistics.

**Extended Data Figure 4:**
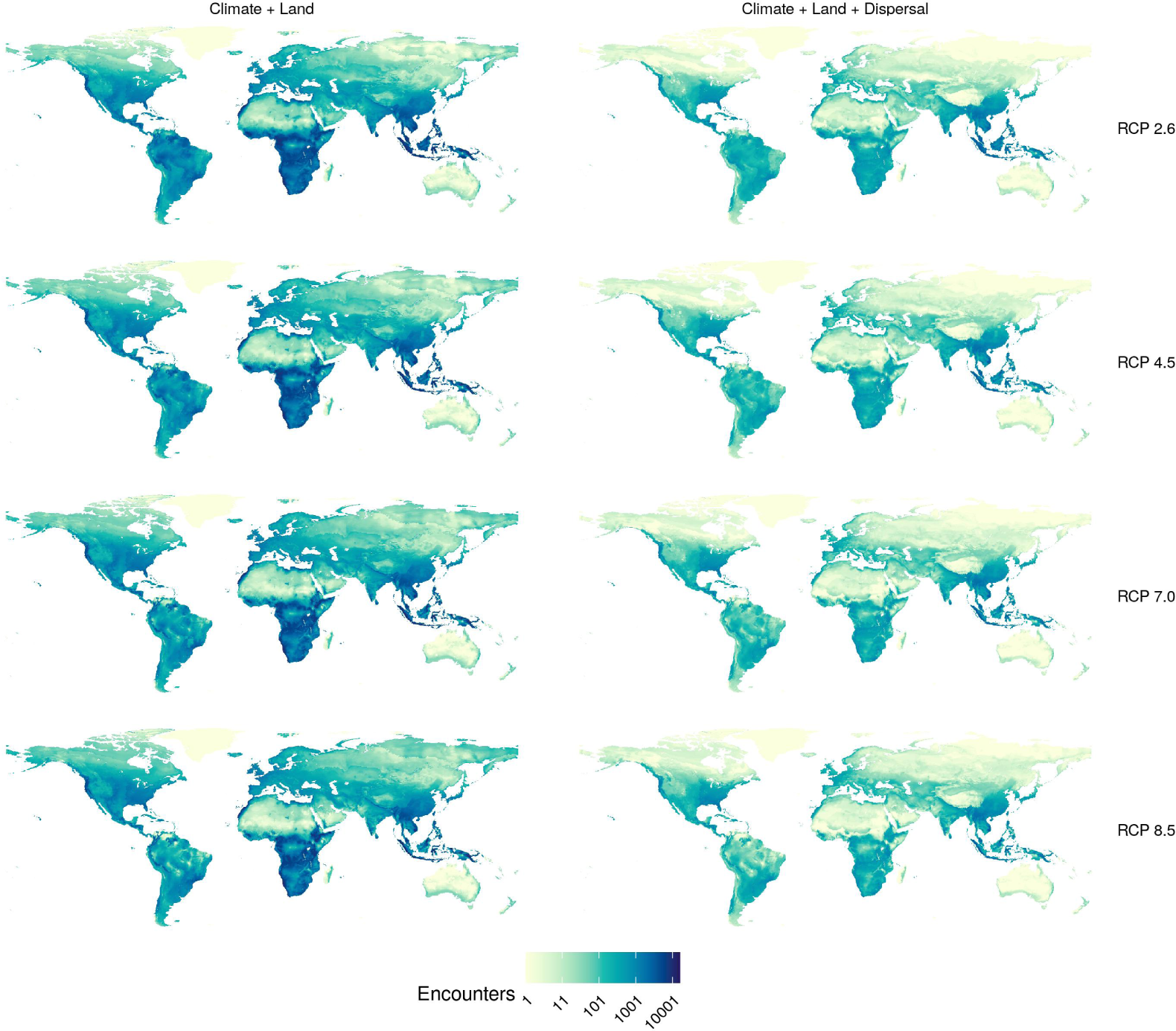
Geographic distribution of first encounters. Predictions were carried out for four representative concentration pathways (RCPs), accounting for climate change and land use change, without (left) and with dispersal limits (right). Darker colours correspond to greater numbers of first encounters in the pixel. Results are averaged across nine global climate models.

**Extended Data Figure 5:**
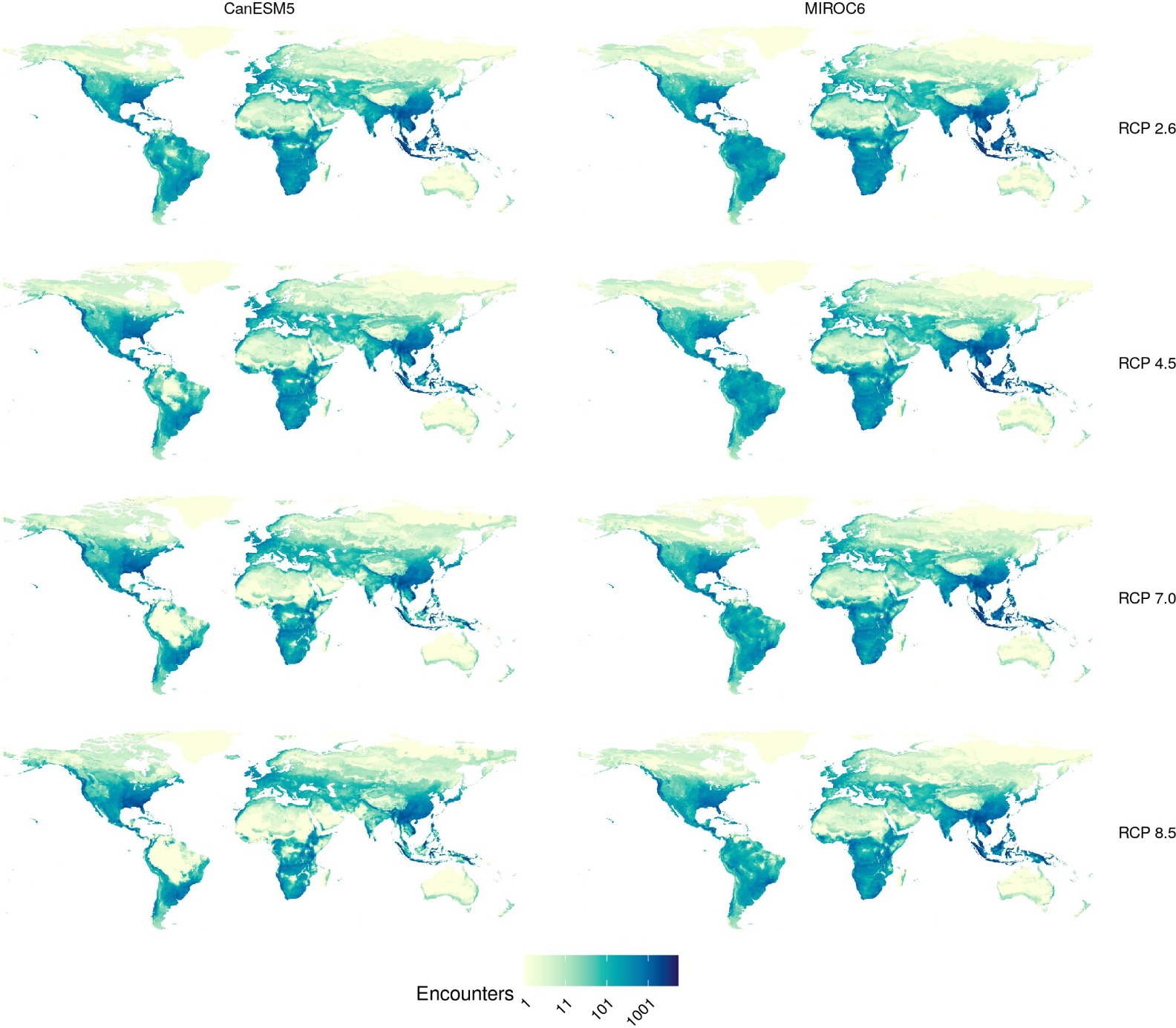
Geographic distribution of first encounters in two global climate models. Predictions were carried out for four representative concentration pathways (RCPs), accounting for climate change and land use change through pairing with shared socioeconomic pathways (SSPs) as detailed in the Methods. The two models selected are those with the highest (CanESM5) and lowest (MIROC6) effective climate sensitivity in the available CMIP6 set on WorldClim ^92^. Darker colours correspond to greater numbers of first encounters in the pixel.

**Extended Data Figure 6:**
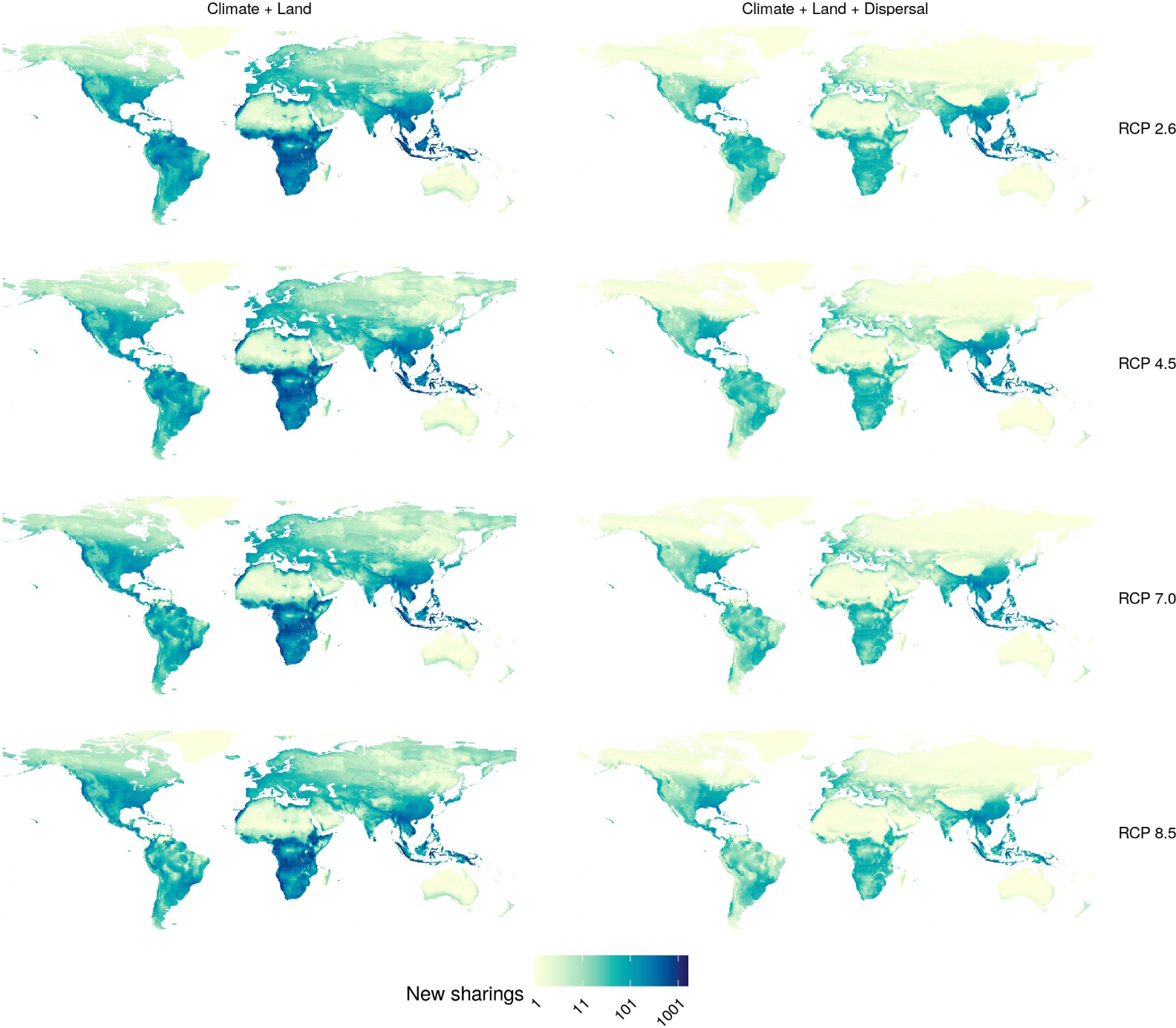
Geographic distribution of expected viral sharing events from first encounters. Predictions were carried out for potential future distributions for four representative concentration pathways (RCPs), accounting for climate change and land use change, without (left) and with dispersal limits (right). Darker colours correspond to greater numbers of new viral sharing events in the pixel. Probability of new viral sharing was calculated by subtracting the species pair ’s present sharing probability from their future sharing probability that our viral sharing GAMMs predicted. This probability was projected across the species pair ’s range intersection, and then summed across all novel species pairs in each pixel. Results are averaged across nine global climate models.

**Extended Data Figure 7:**
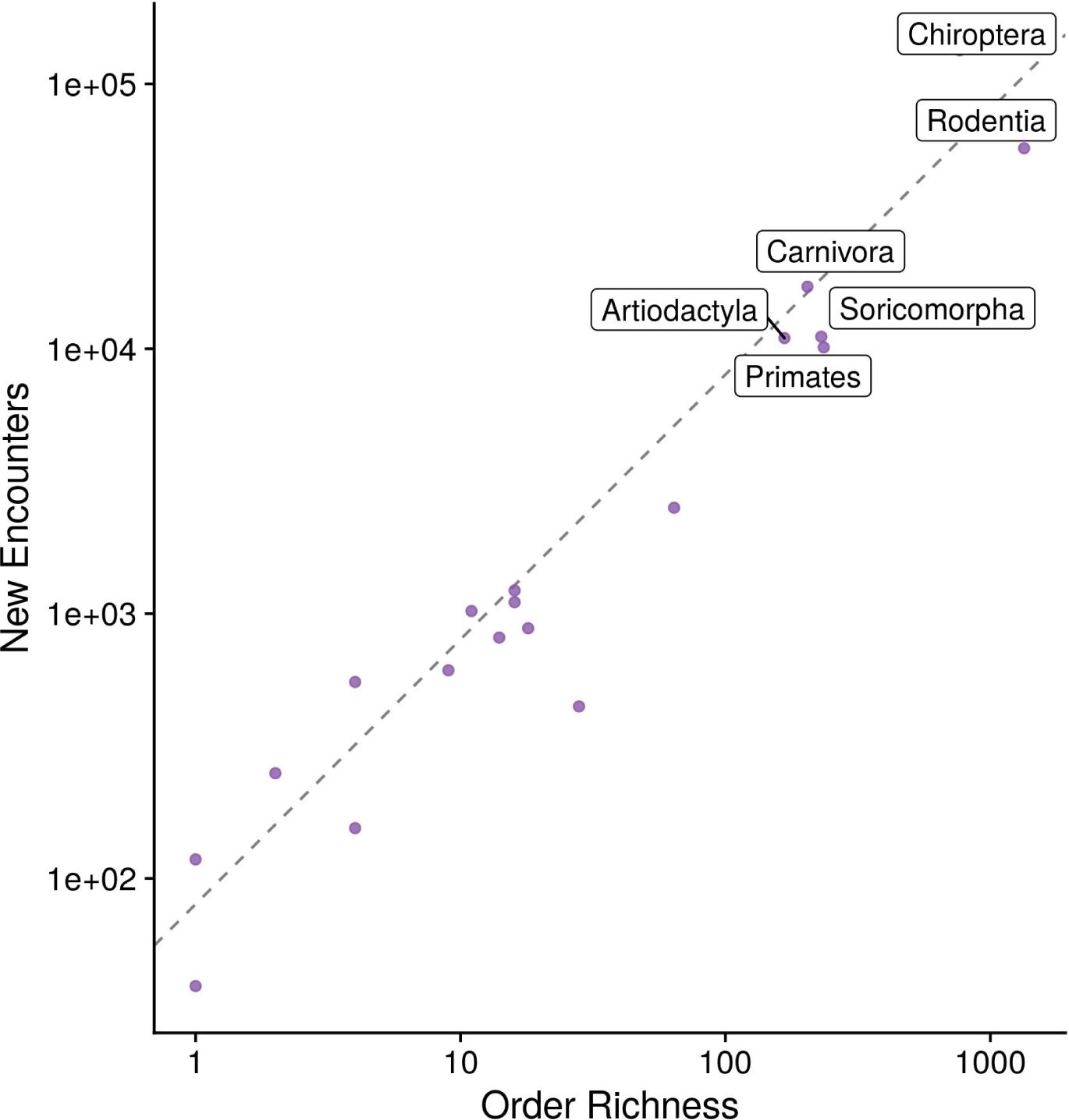
Order-level heterogeneity in first encounters. Dispersal stratifies the number of first encounters (RCP 2.6 with all range filters), where some orders have more than expected at random, based on the mean number of first encounters and order size (line). Results are averaged across nine global climate models.

**Extended Data Figure 8:**
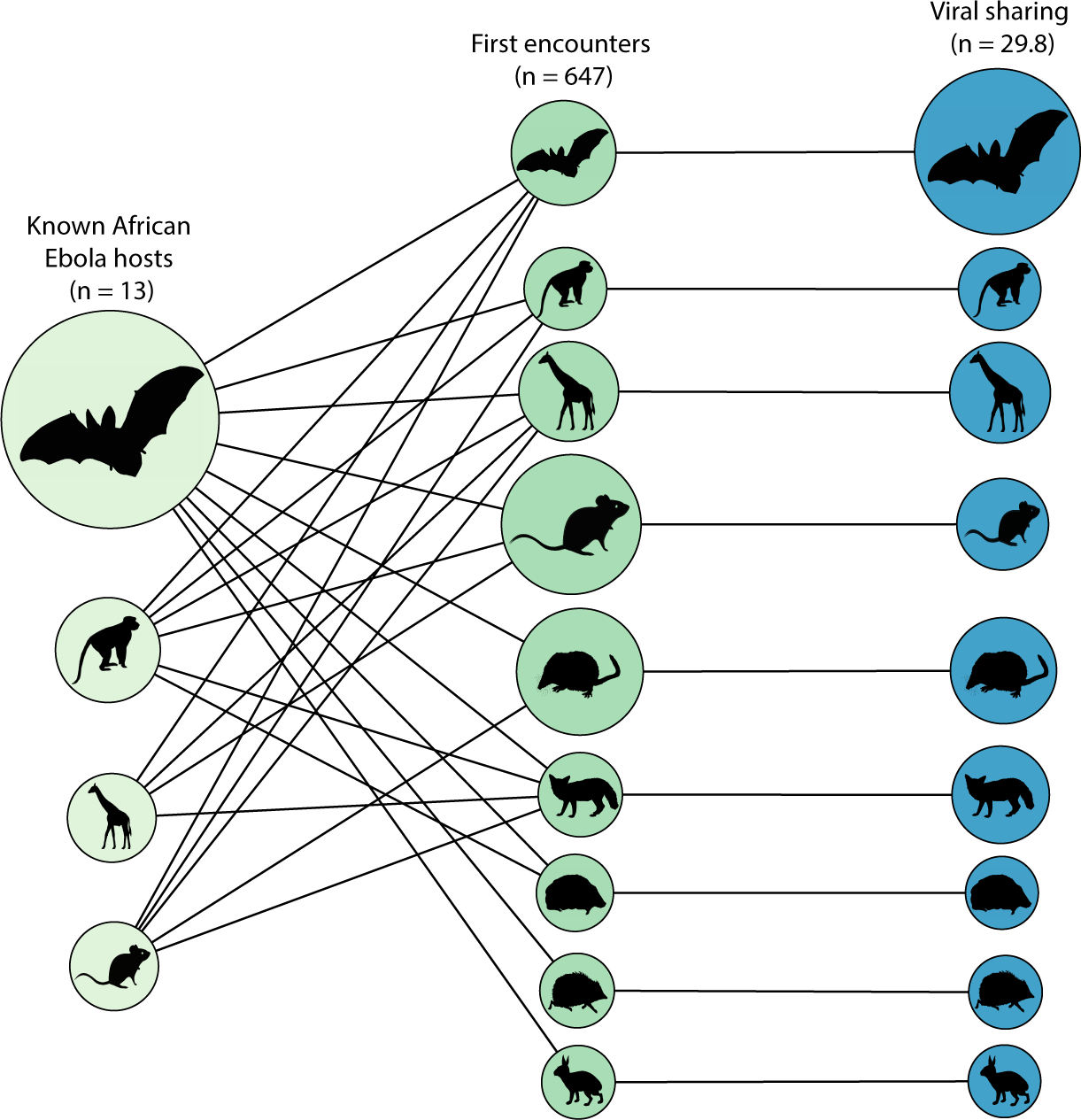
Projected viral sharing from suspected Ebola reservoirs is dominated by bats. Node size is proportional to (left) the number of suspected Ebola host species in each order, which connect to (middle) first encounters with potentially naive host species; and (right) the number of projected viral sharing events in each receiving group. (Node size denotes proportions out of 100% within each column total.) While Ebola hosts will encounter a much wider taxonomic range of mammal groups than current reservoirs, the vast majority of future viral sharing will occur disproportionately in bats.

**Extended Data Figure 9:**
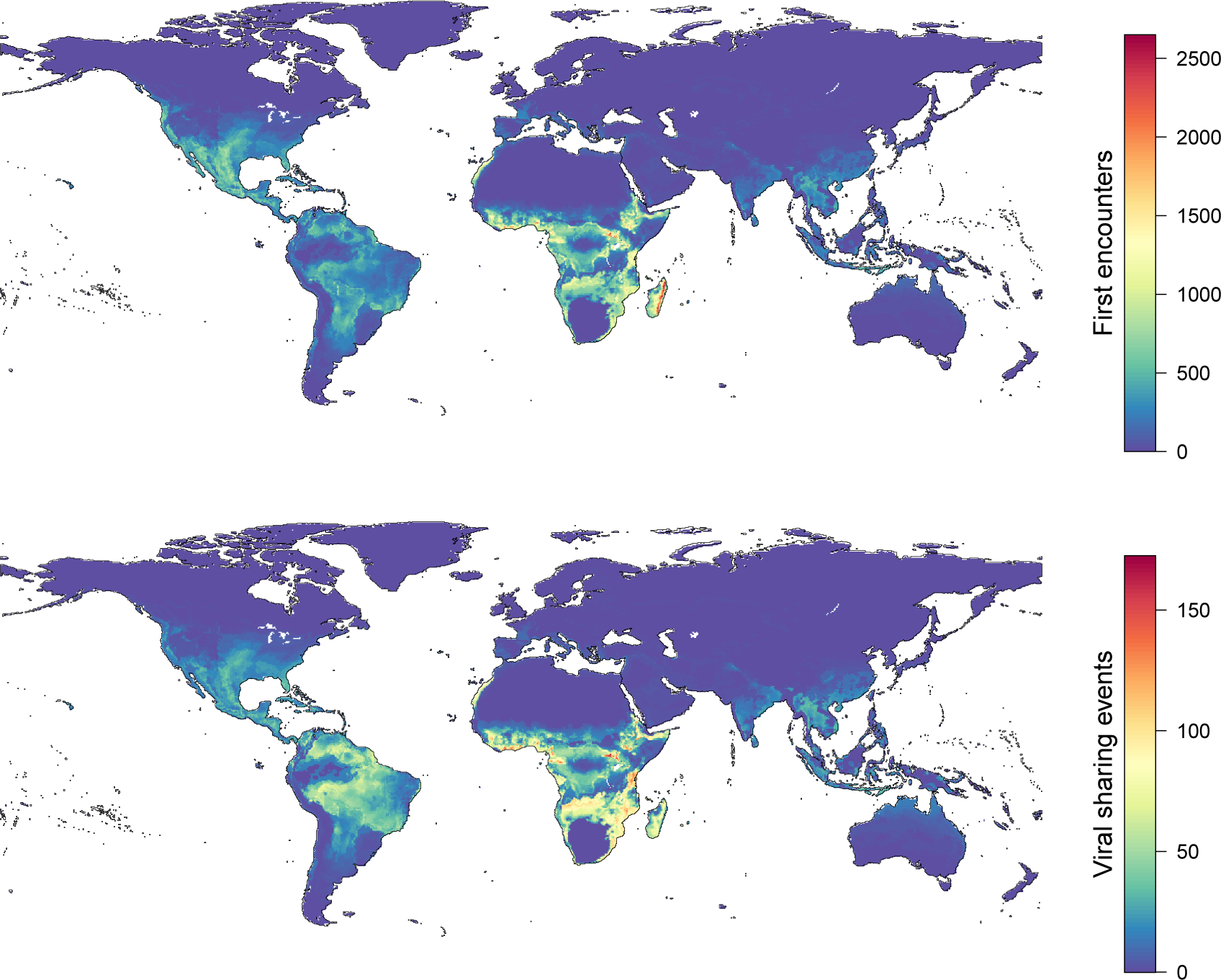
Projected viral sharing from present-day warming. First encounters and viral sharing events are derived from an independent analysis of ERA5 climate data for the present day (2005-2019) versus the recent past (1981-1995).

**Extended Data Figure 10:**
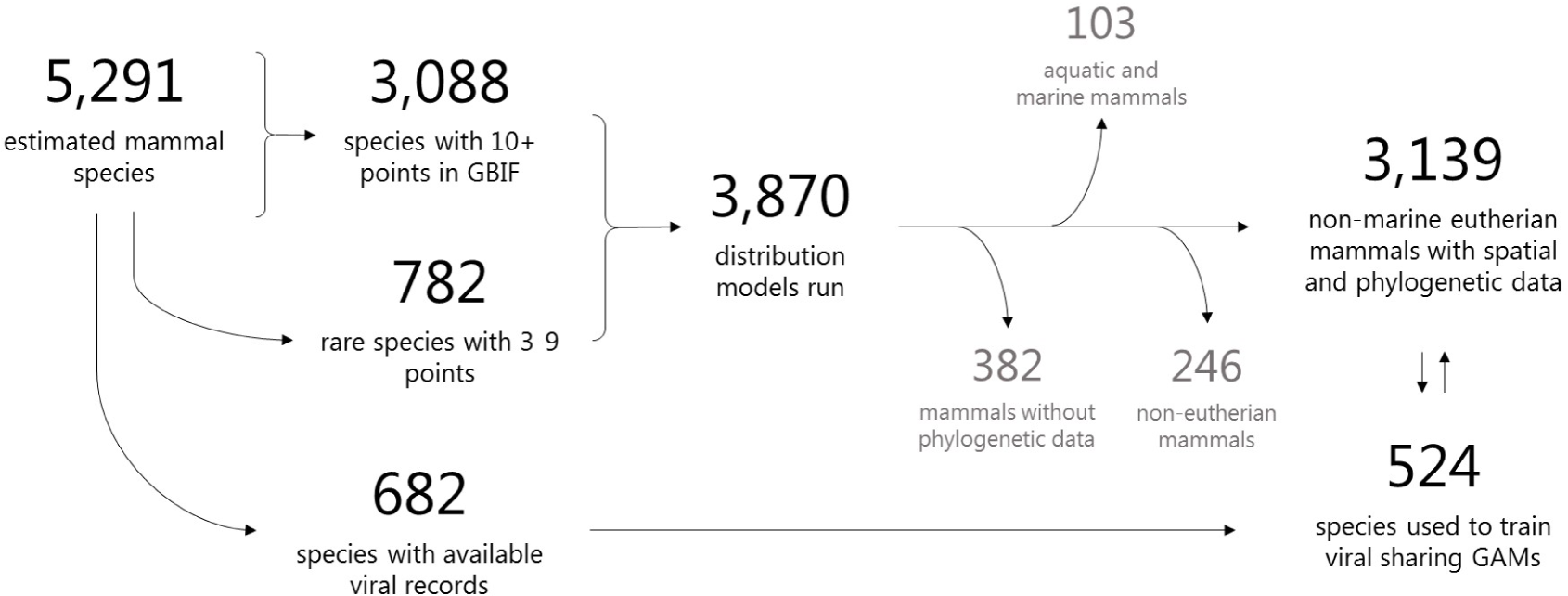
Data processing workflow. Summary of species inclusion across the modeling pipeline for species distributions and viral sharing models. The final analyses in the main text use 3,139 species of placental mammals across all scenarios.

**Extended Data Figure 11:**
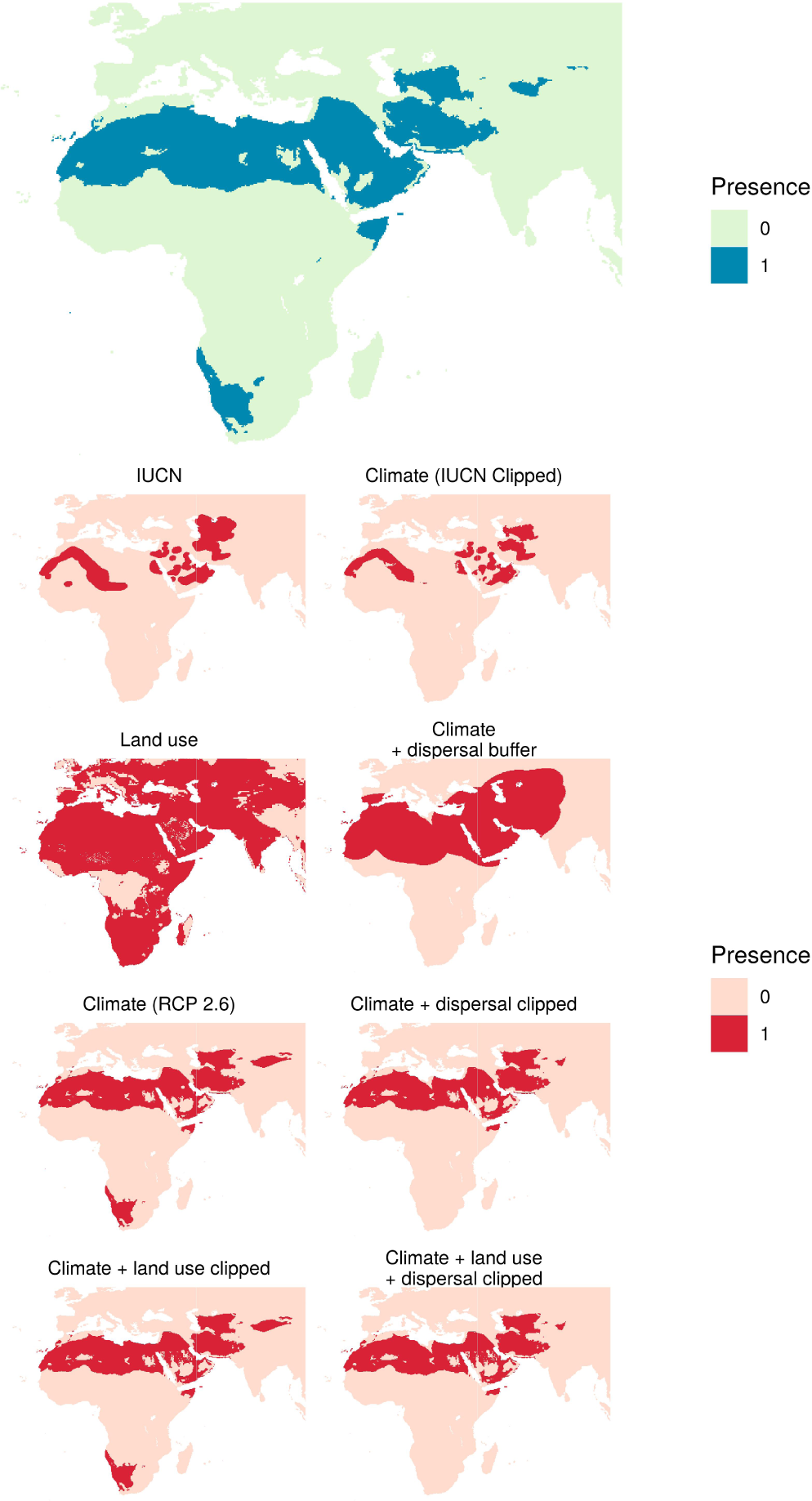
Species distribution modeling workflow for a single species. A focal species (the sand cat, *Felis margarita*) is displayed as an illustrative example. The present day climate prediction (top left) was clipped to the same continent according to the IUCN distribution (top right). This was then clipped according to *Felis margarita*’s land use (second row, left). The known dispersal distance of the sand cat was used to buffer the climate distribution (second row, right). The potential future distribution predictions (RCP 2.6 shown as an example) are displayed in the bottom four panels, for each of the four pipelines: only climate (third row, left); climate + dispersal clip (third row, right); climate + land use clip (bottom row, left) and climate + land use + dispersal clip (bottom row, right). The four distributions clearly display the limiting effect of the dispersal filter (bottom right panels) in reducing the probability of novel species interactions (bottom left panels). The land use clip had little effect on this species as the entire distribution area was habitable for the sand cat.

