## Supplemental Figures for "Climate change will drive novel cross-species viral transmission"

**Figure S1. Geographic distribution of first encounters in BCC-CSM2-MR.** Predictions were carried out for four representative concentration pathways (RCPs), accounting for climate change and land use change, without (left) and with dispersal limits (right). Darker colours correspond to greater numbers of first encounters in the pixel.

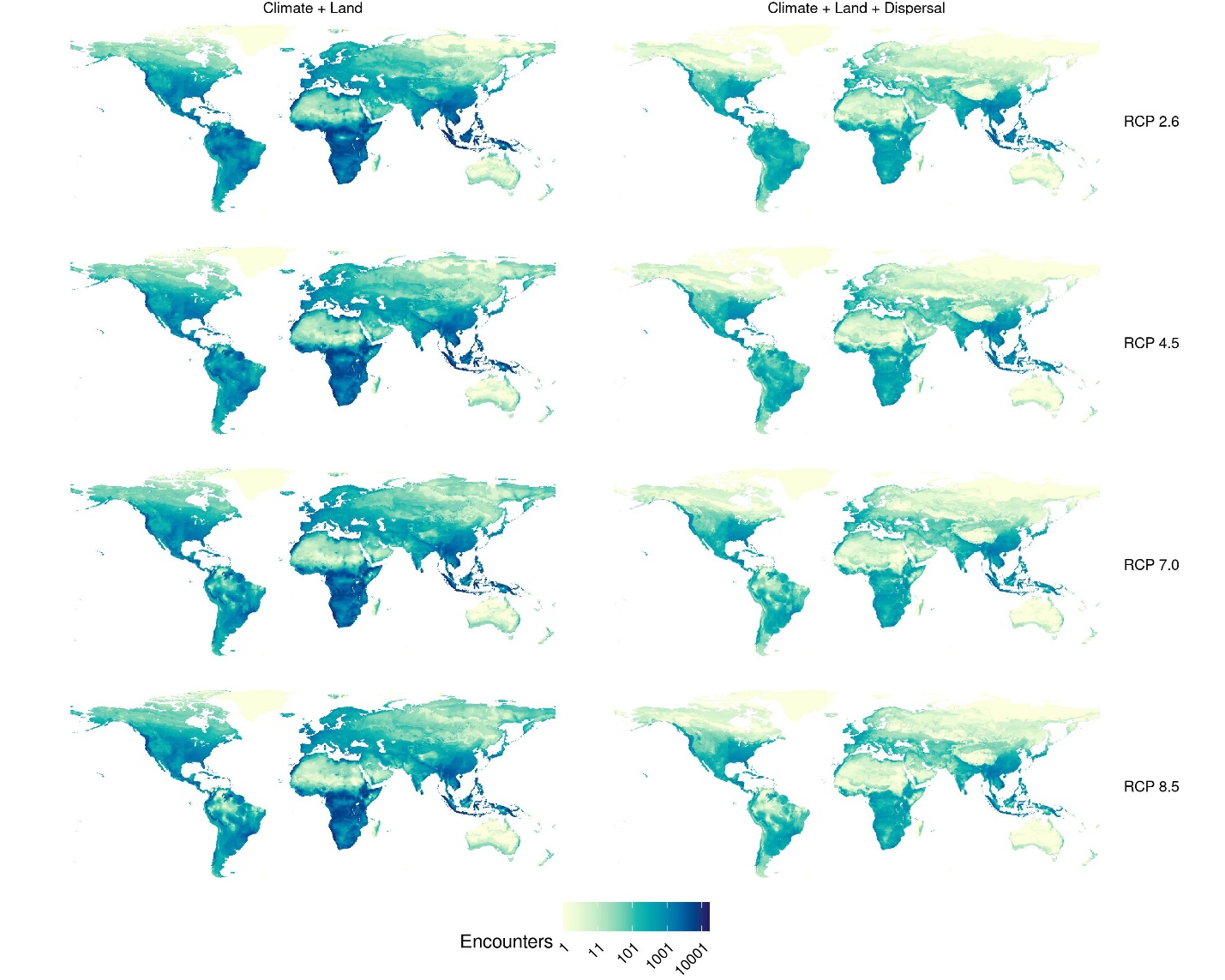

**Figure S2. Geographic distribution of first encounters in CanESM5.** Predictions were carried out for four representative concentration pathways (RCPs), accounting for climate change and land use change, without (left) and with dispersal limits (right). Darker colours correspond to greater numbers of first encounters in the pixel.

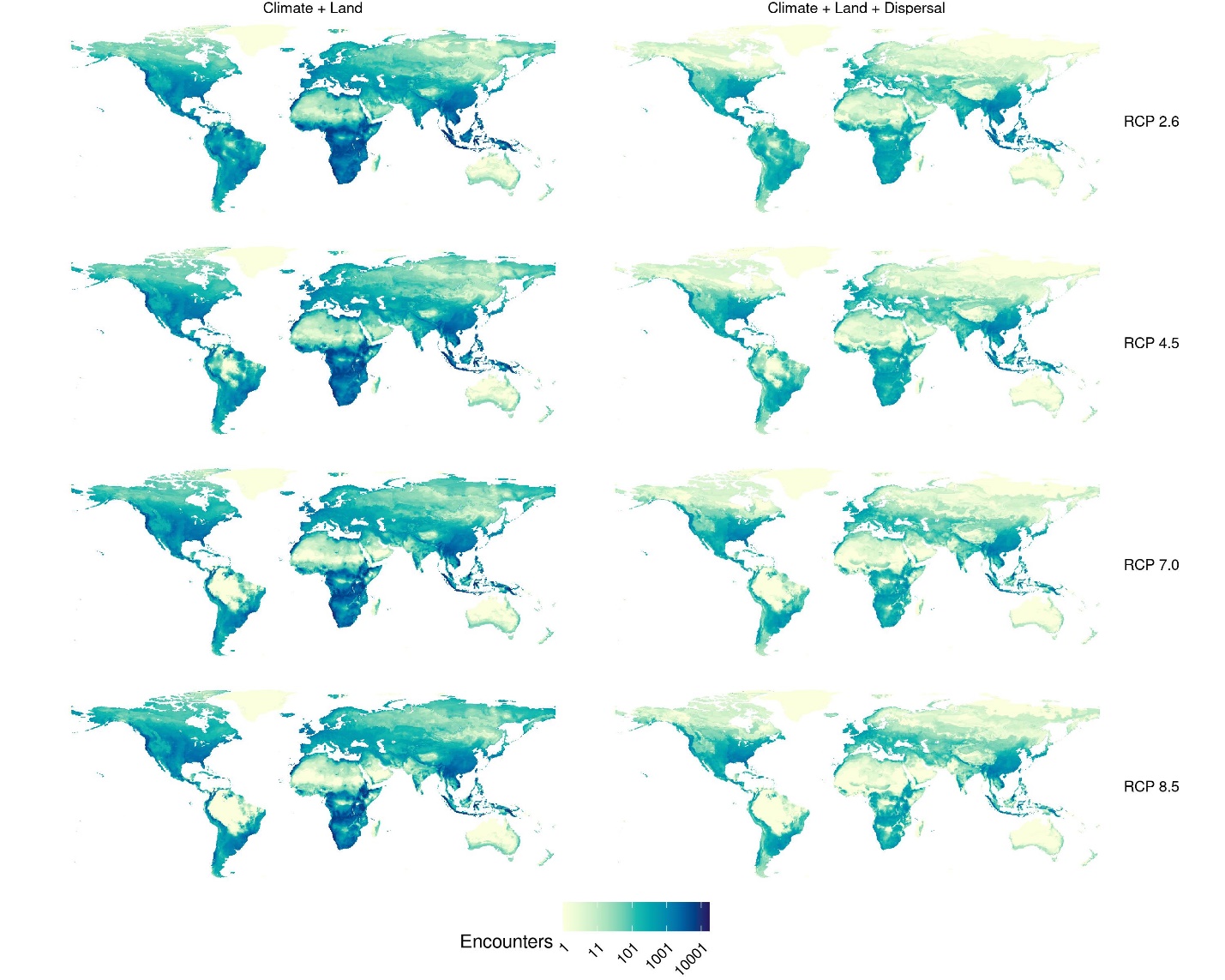

**Figure S3. Geographic distribution of first encounters in CNRM-CM6-1.** Predictions were carried out for four representative concentration pathways (RCPs), accounting for climate change and land use change, without (left) and with dispersal limits (right). Darker colours correspond to greater numbers of first encounters in the pixel.

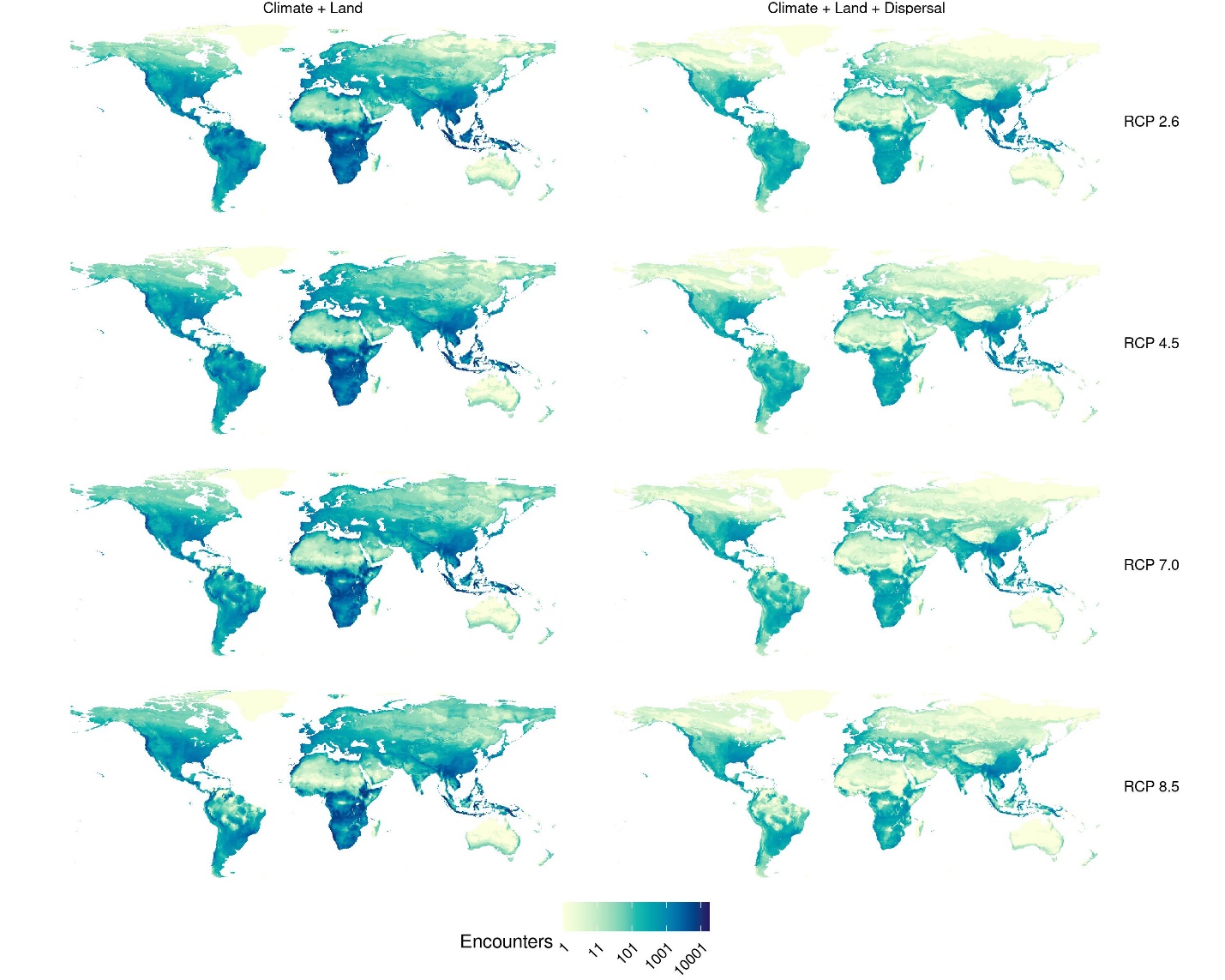

**Figure S4. Geographic distribution of first encounters in CNRM-ESM2-1.** Predictions were carried out for four representative concentration pathways (RCPs), accounting for climate change and land use change, without (left) and with dispersal limits (right). Darker colours correspond to greater numbers of first encounters in the pixel.

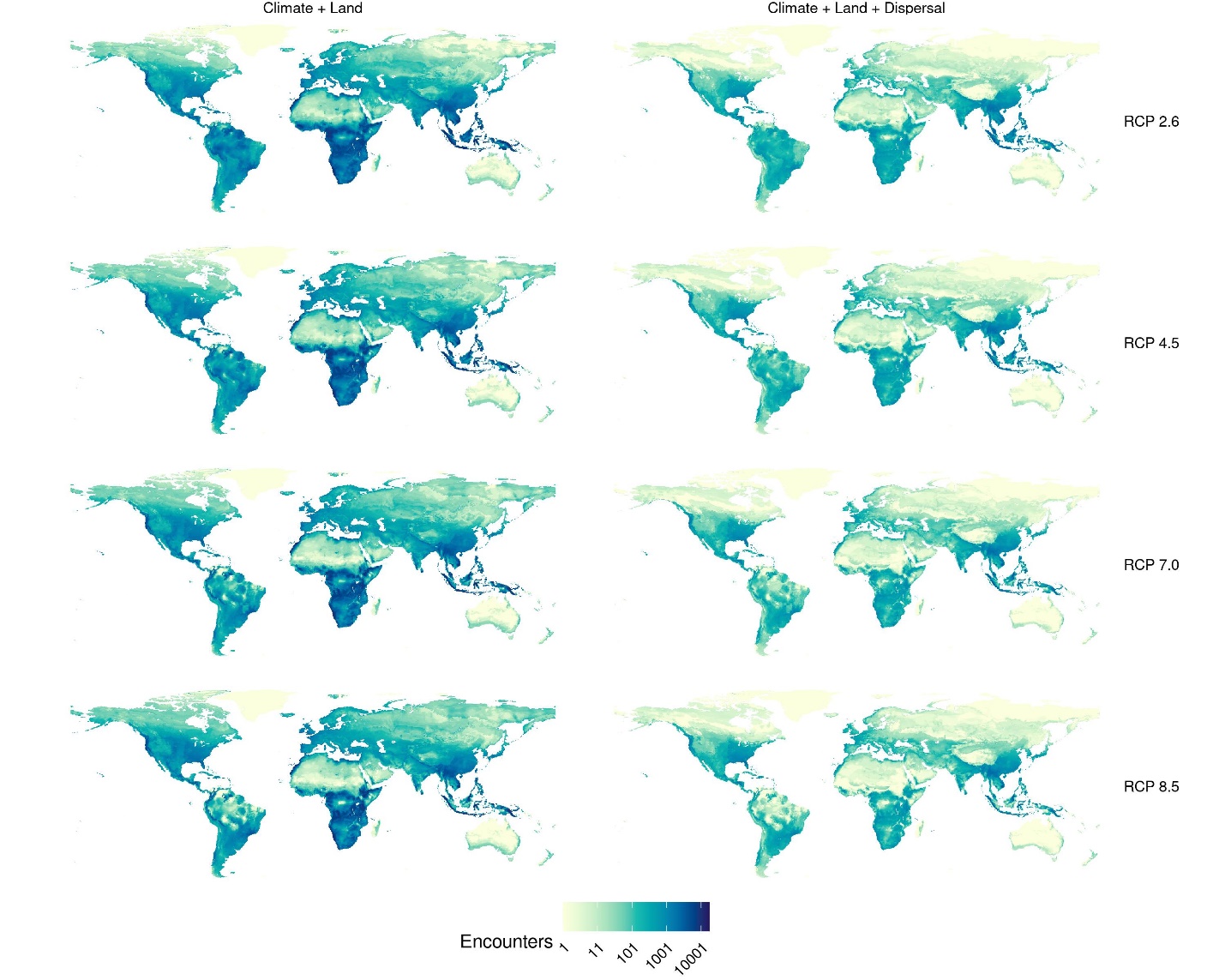

**Figure S5. Geographic distribution of first encounters in GFDL-ESM4.** Predictions were carried out for the only two available representative concentration pathways (RCPs; see methods), accounting for climate change and land use change, without (left) and with dispersal limits (right). Darker colours correspond to greater numbers of first encounters in the pixel.

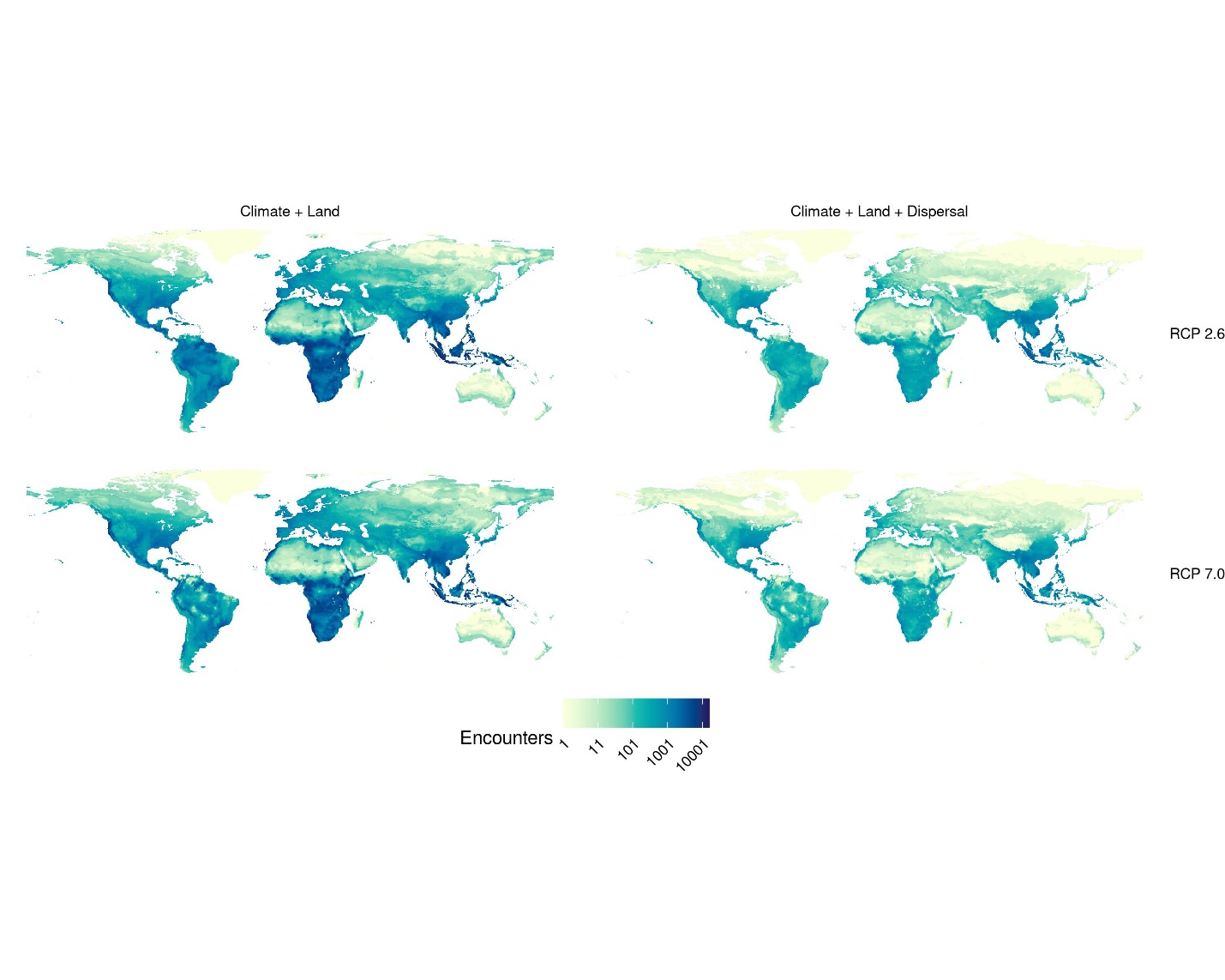

**Figure S6. Geographic distribution of first encounters in IPSL-CM6A-LR.** Predictions were carried out for four representative concentration pathways (RCPs), accounting for climate change and land use change, without (left) and with dispersal limits (right). Darker colours correspond to greater numbers of first encounters in the pixel.

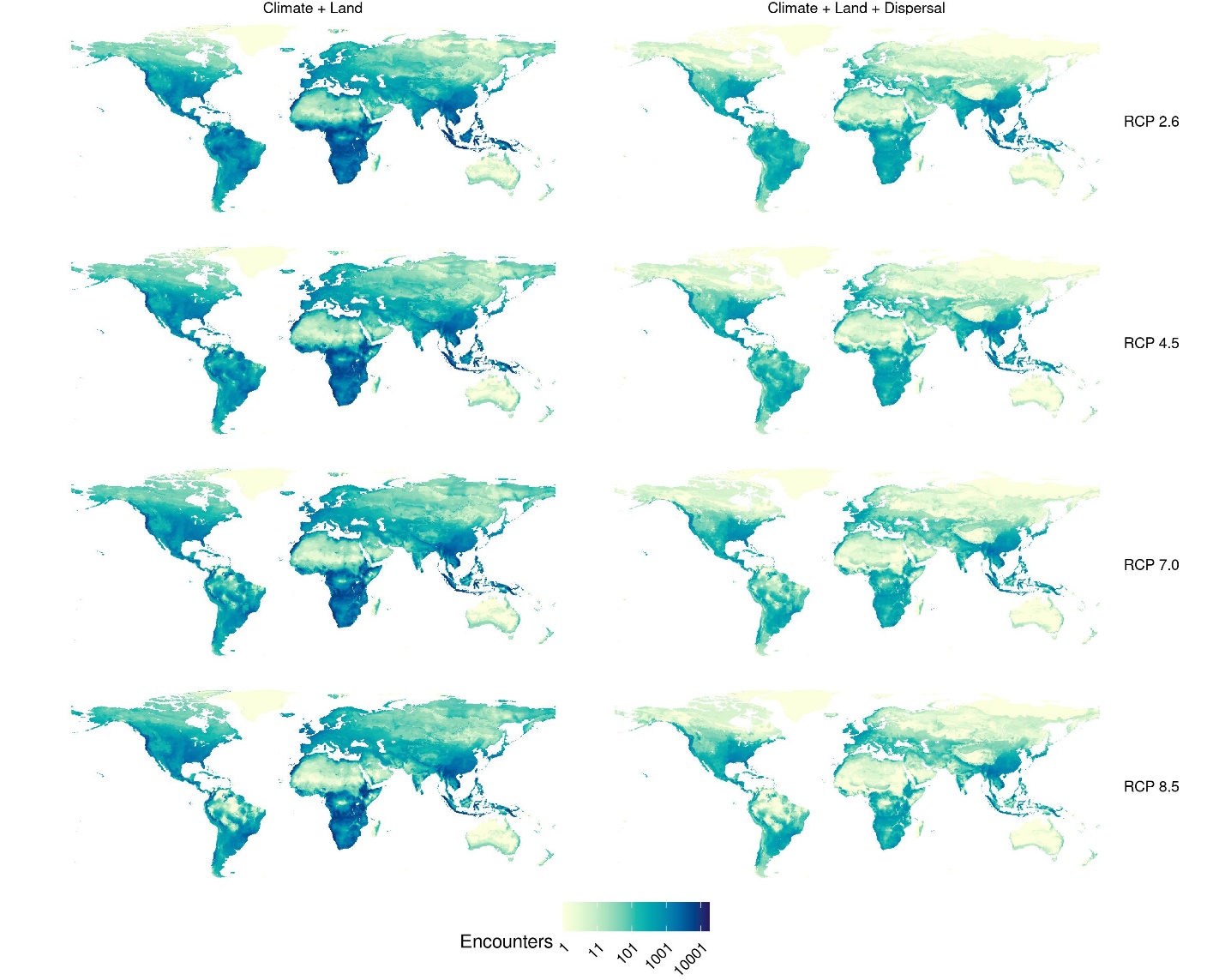

**Figure S7. Geographic distribution of first encounters in MIROC-ES2L.** Predictions were carried out for four representative concentration pathways (RCPs), accounting for climate change and land use change, without (left) and with dispersal limits (right). Darker colours correspond to greater numbers of first encounters in the pixel.

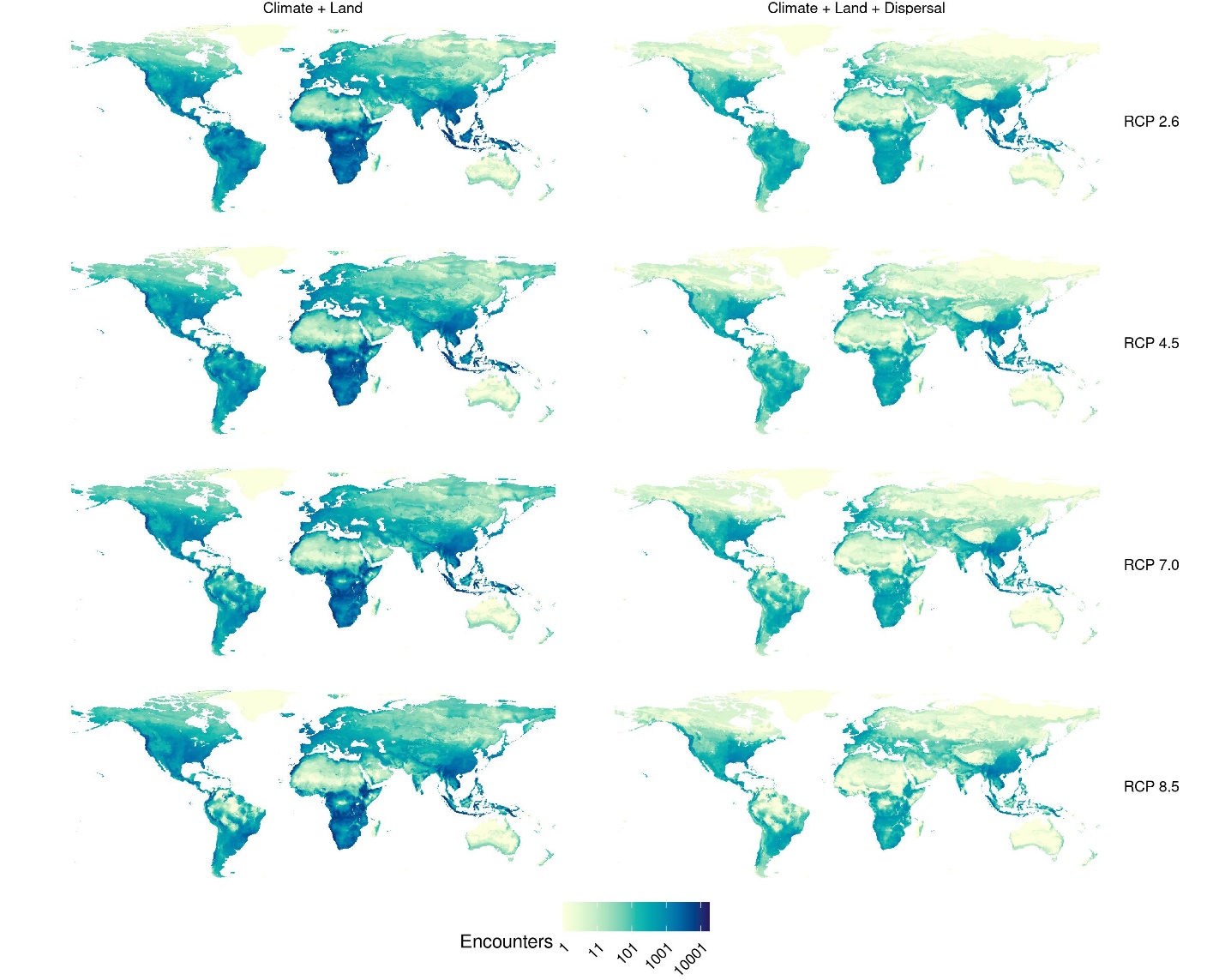

**Figure S8. Geographic distribution of first encounters in MIROC6.** Predictions were carried out for four representative concentration pathways (RCPs), accounting for climate change and land use change, without (left) and with dispersal limits (right). Darker colours correspond to greater numbers of first encounters in the pixel.

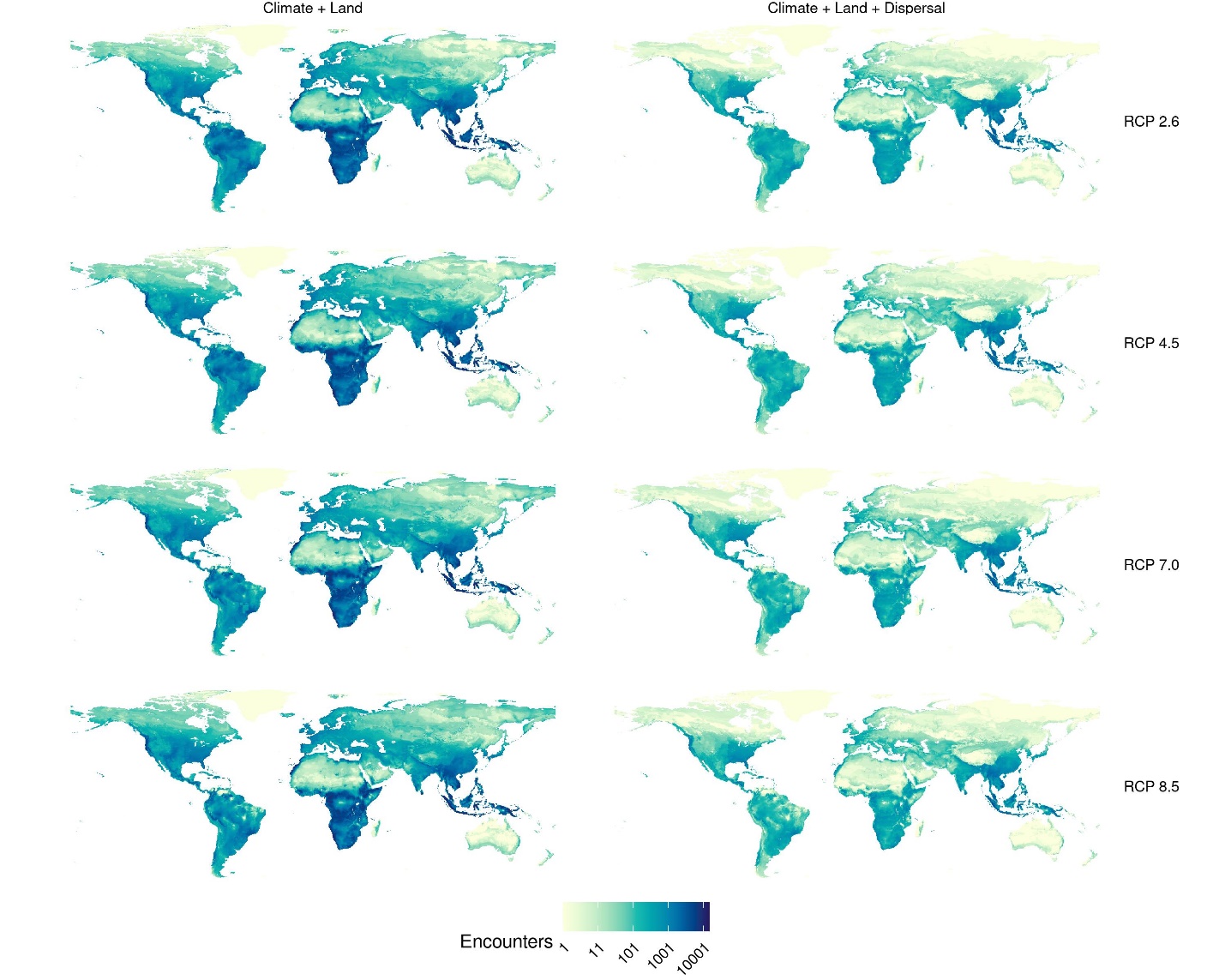

**Figure S9. Geographic distribution of first encounters in MRI-ESM2-0.** Predictions were carried out for four representative concentration pathways (RCPs), accounting for climate change and land use change, without (left) and with dispersal limits (right). Darker colours correspond to greater numbers of first encounters in the pixel.

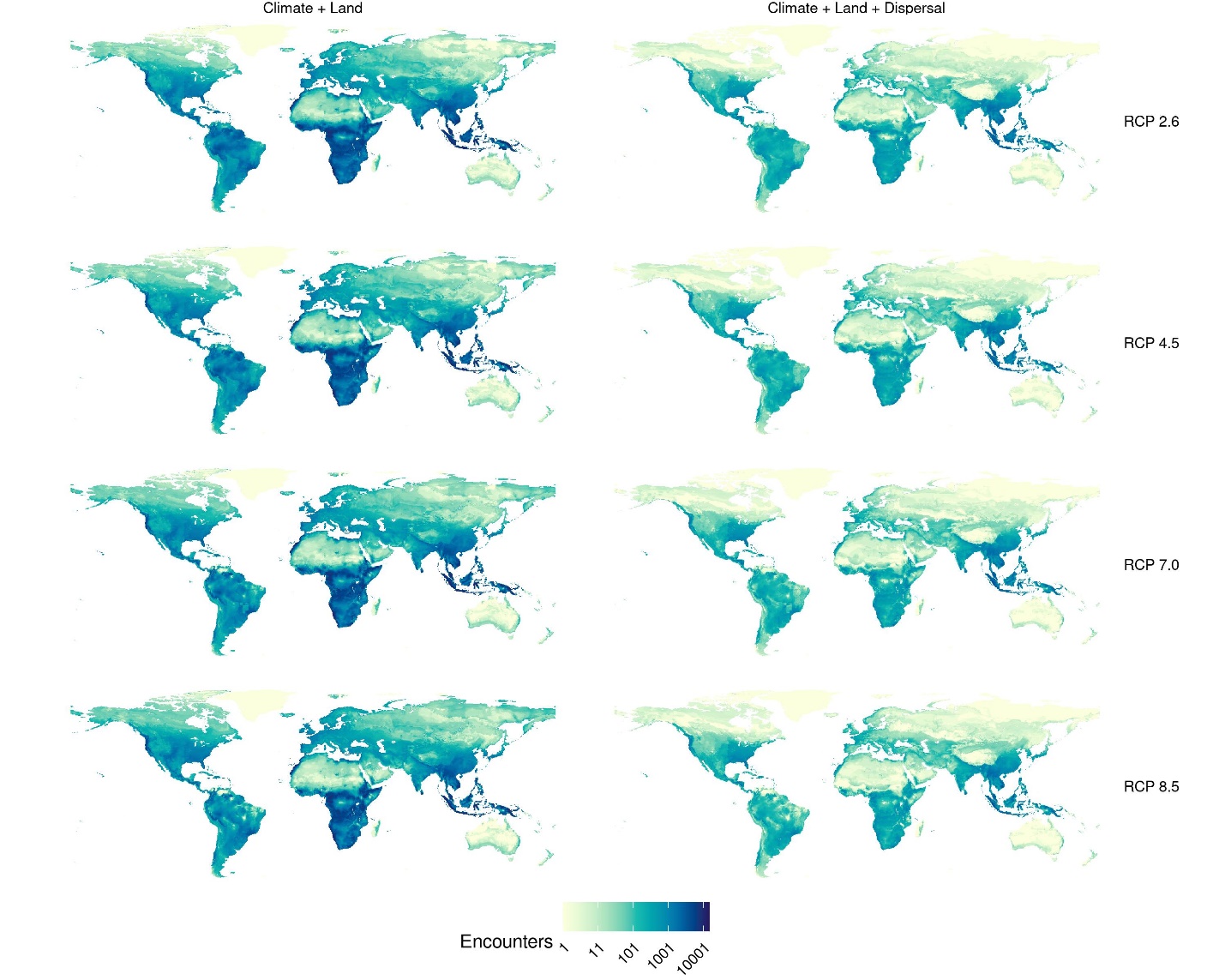

**Figure S10. Geographic distribution of viral sharing events in BCC-CSM2-MR.** Predictions were carried out for four representative concentration pathways (RCPs), accounting for climate change and land use change, without (left) and with dispersal limits (right). Darker colours correspond to greater numbers of viral sharing events in the pixel.

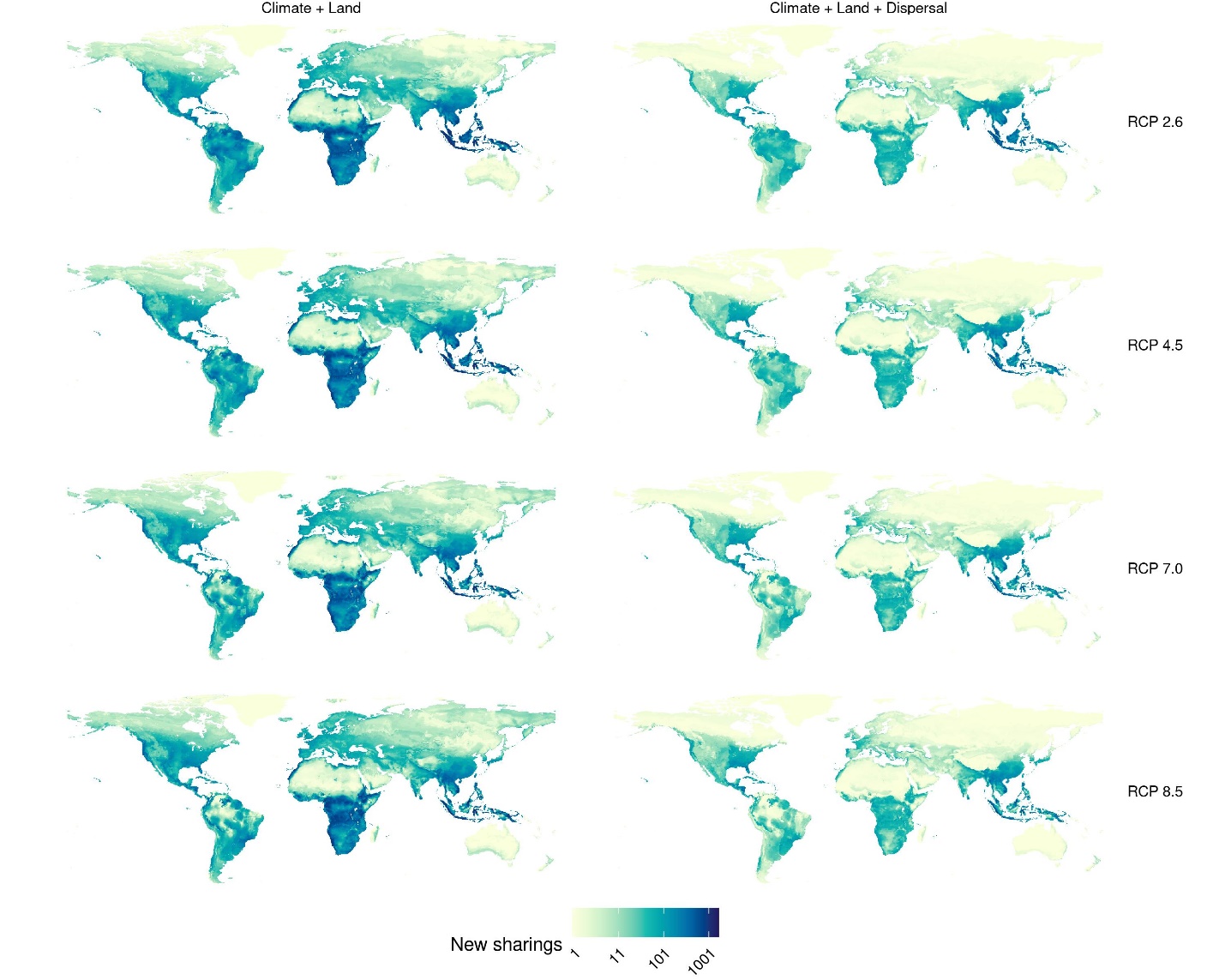

**Figure S11. Geographic distribution of viral sharing events in CanESM5.** Predictions were carried out for four representative concentration pathways (RCPs), accounting for climate change and land use change, without (left) and with dispersal limits (right). Darker colours correspond to greater numbers of viral sharing events in the pixel.

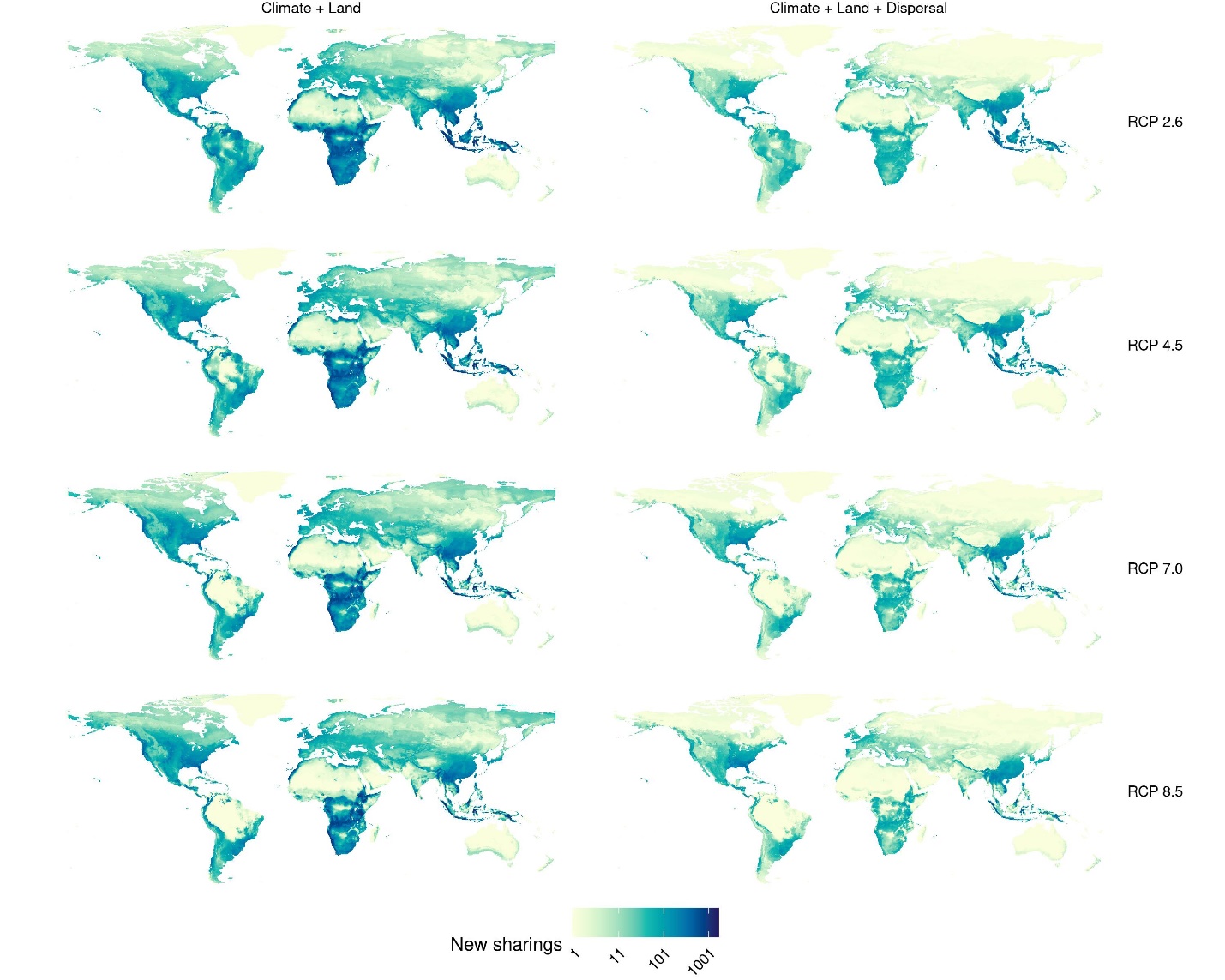

**Figure S12. Geographic distribution of viral sharing events in CNRM-CM6-1.** Predictions were carried out for four representative concentration pathways (RCPs), accounting for climate change and land use change, without (left) and with dispersal limits (right). Darker colours correspond to greater numbers of viral sharing events in the pixel.

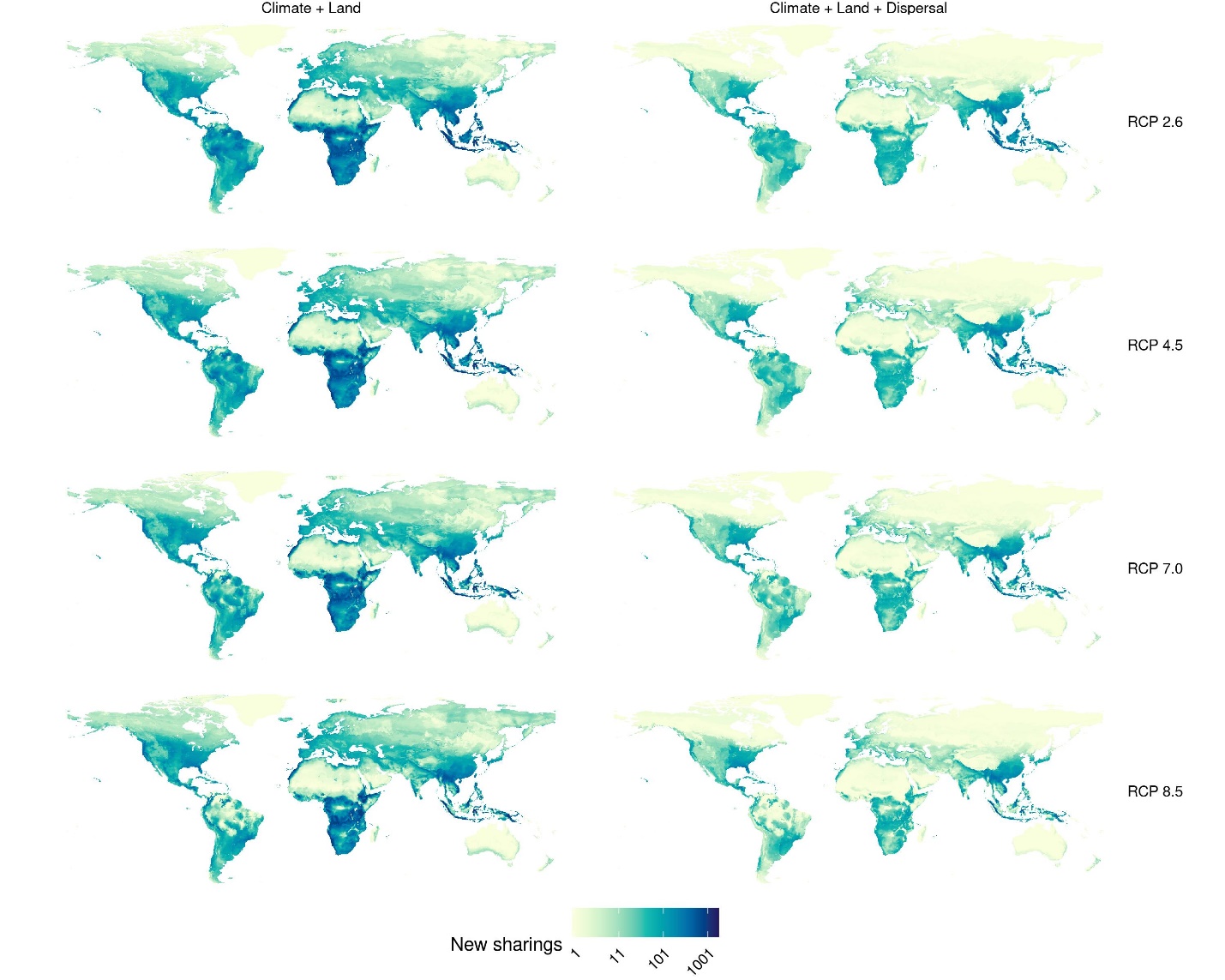

**Figure S13. Geographic distribution of viral sharing events in CNRM-ESM2-1.** Predictions were carried out for four representative concentration pathways (RCPs), accounting for climate change and land use change, without (left) and with dispersal limits (right). Darker colours correspond to greater numbers of viral sharing events in the pixel.

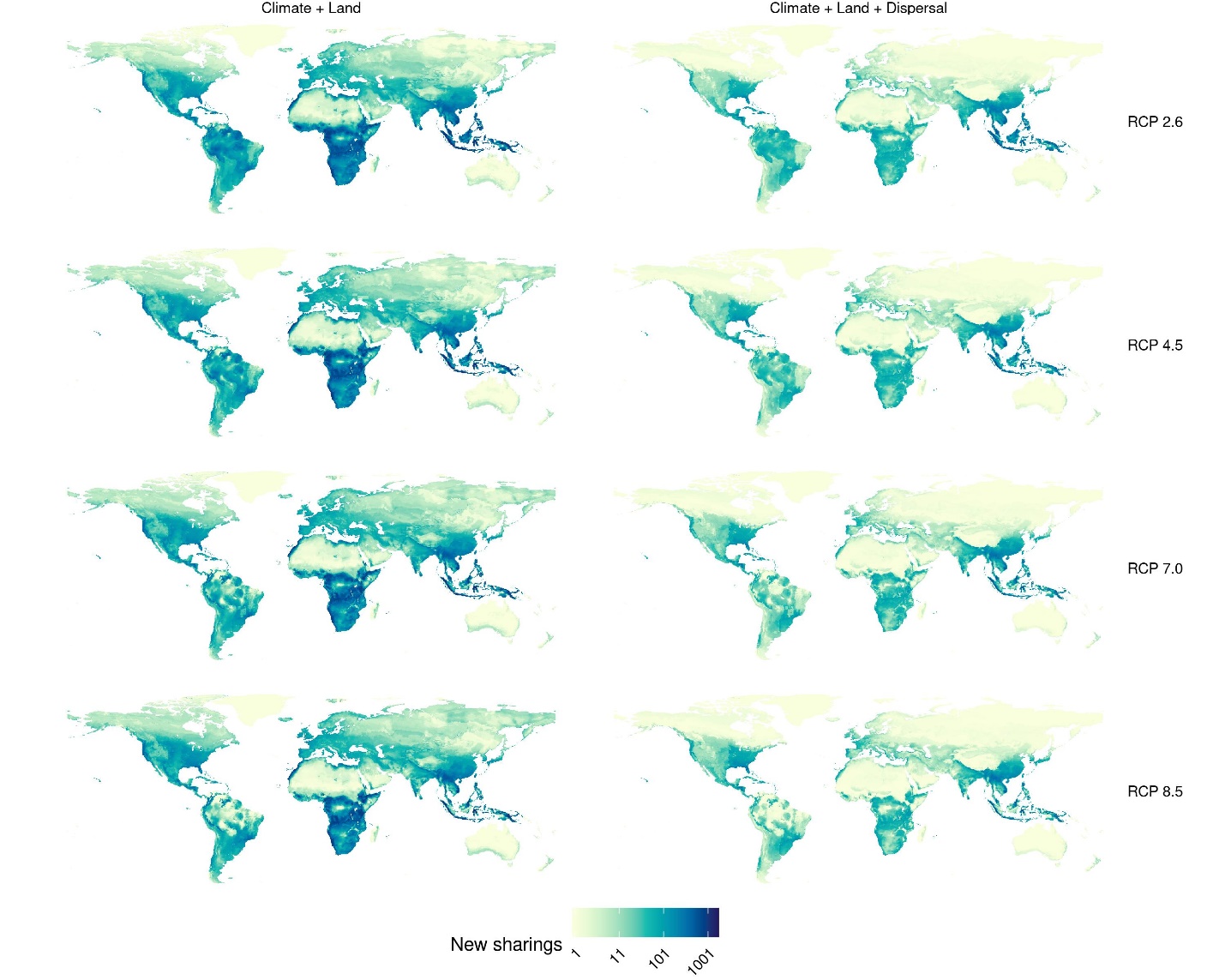

**Figure S14. Geographic distribution of viral sharing events in GFDL-ESM4.** Predictions were carried out for the only two available representative concentration pathways (RCPs; see methods), accounting for climate change and land use change, without (left) and with dispersal limits (right). Darker colours correspond to greater numbers of viral sharing events in the pixel.

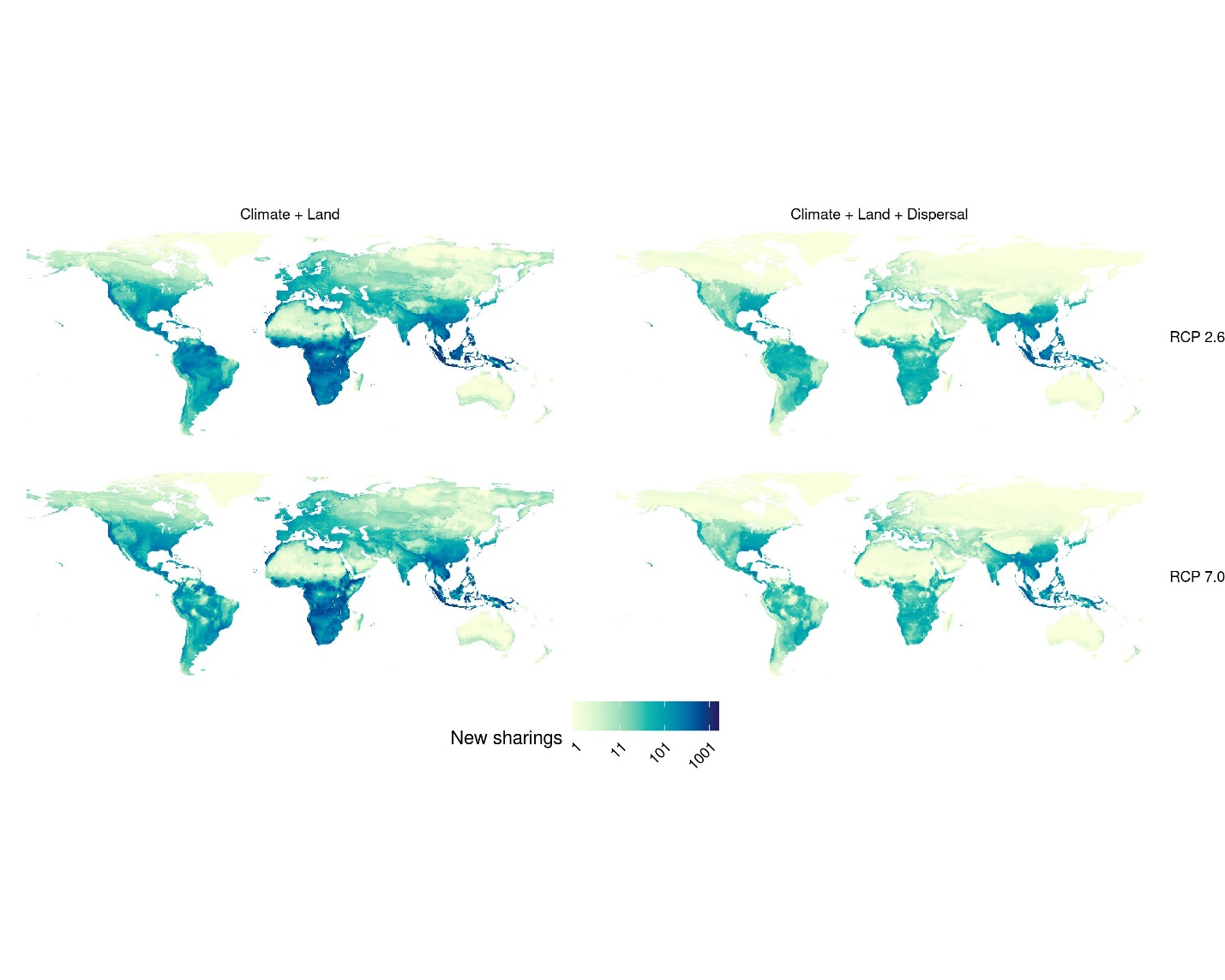

**Figure S15. Geographic distribution of viral sharing events in IPSL-CM6A-LR.** Predictions were carried out for four representative concentration pathways (RCPs), accounting for climate change and land use change, without (left) and with dispersal limits (right). Darker colours correspond to greater numbers of viral sharing events in the pixel.

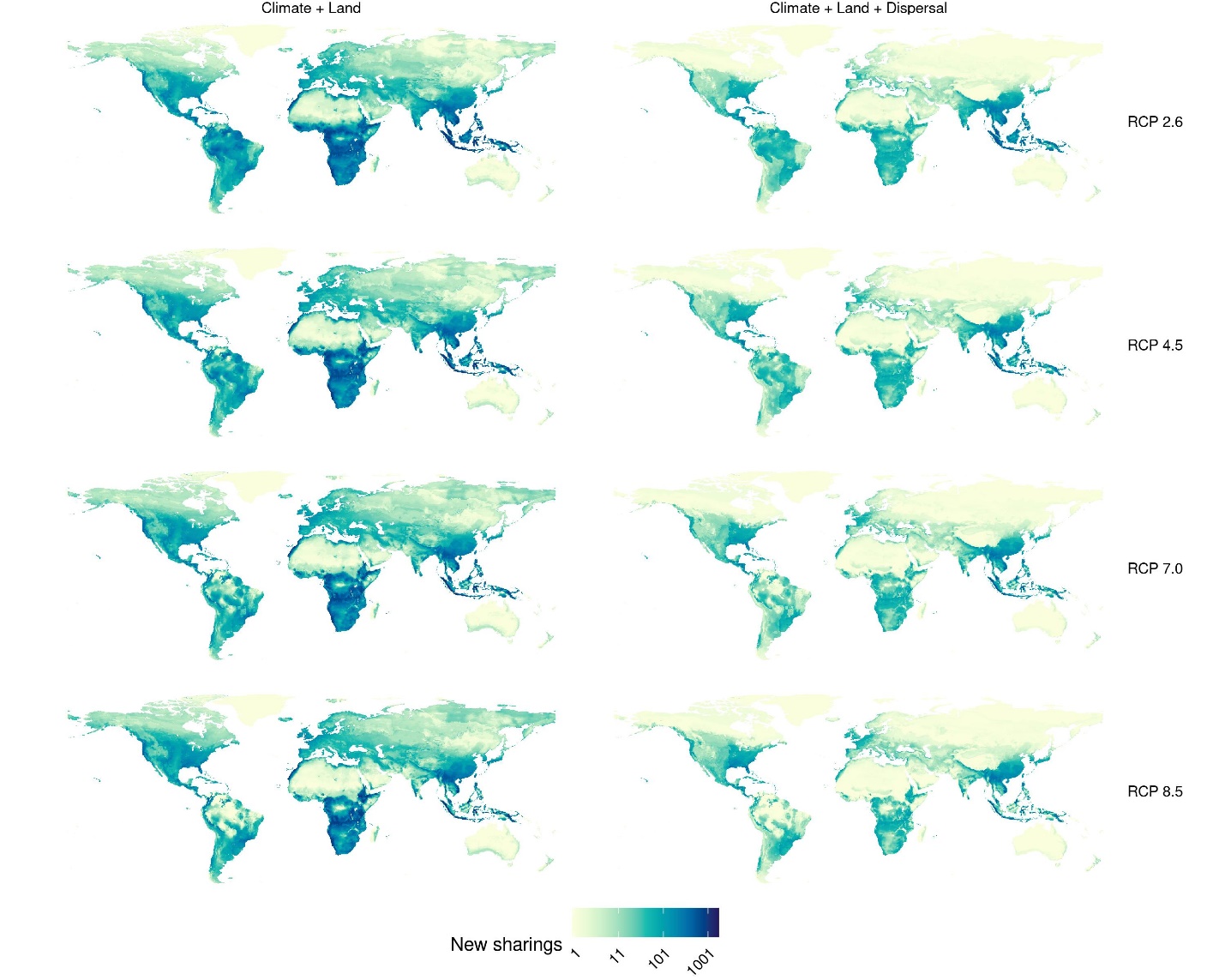

**Figure S16. Geographic distribution of viral sharing events in MIROC-ES2L.** Predictions were carried out for four representative concentration pathways (RCPs), accounting for climate change and land use change, without (left) and with dispersal limits (right). Darker colours correspond to greater numbers of viral sharing events in the pixel.

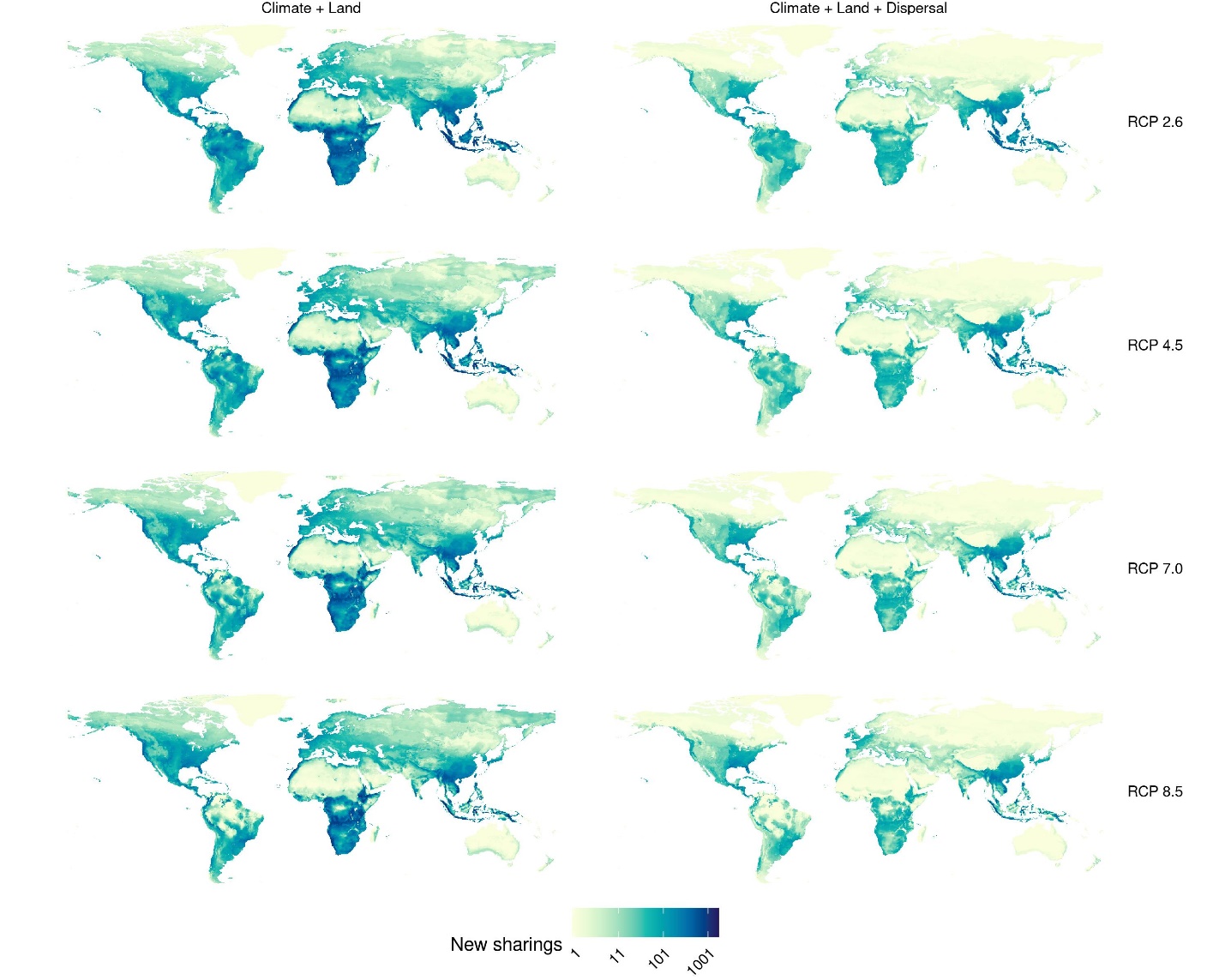

**Figure S17. Geographic distribution of viral sharing events in MIROC6.** Predictions were carried out for four representative concentration pathways (RCPs), accounting for climate change and land use change, without (left) and with dispersal limits (right). Darker colours correspond to greater numbers of viral sharing events in the pixel.

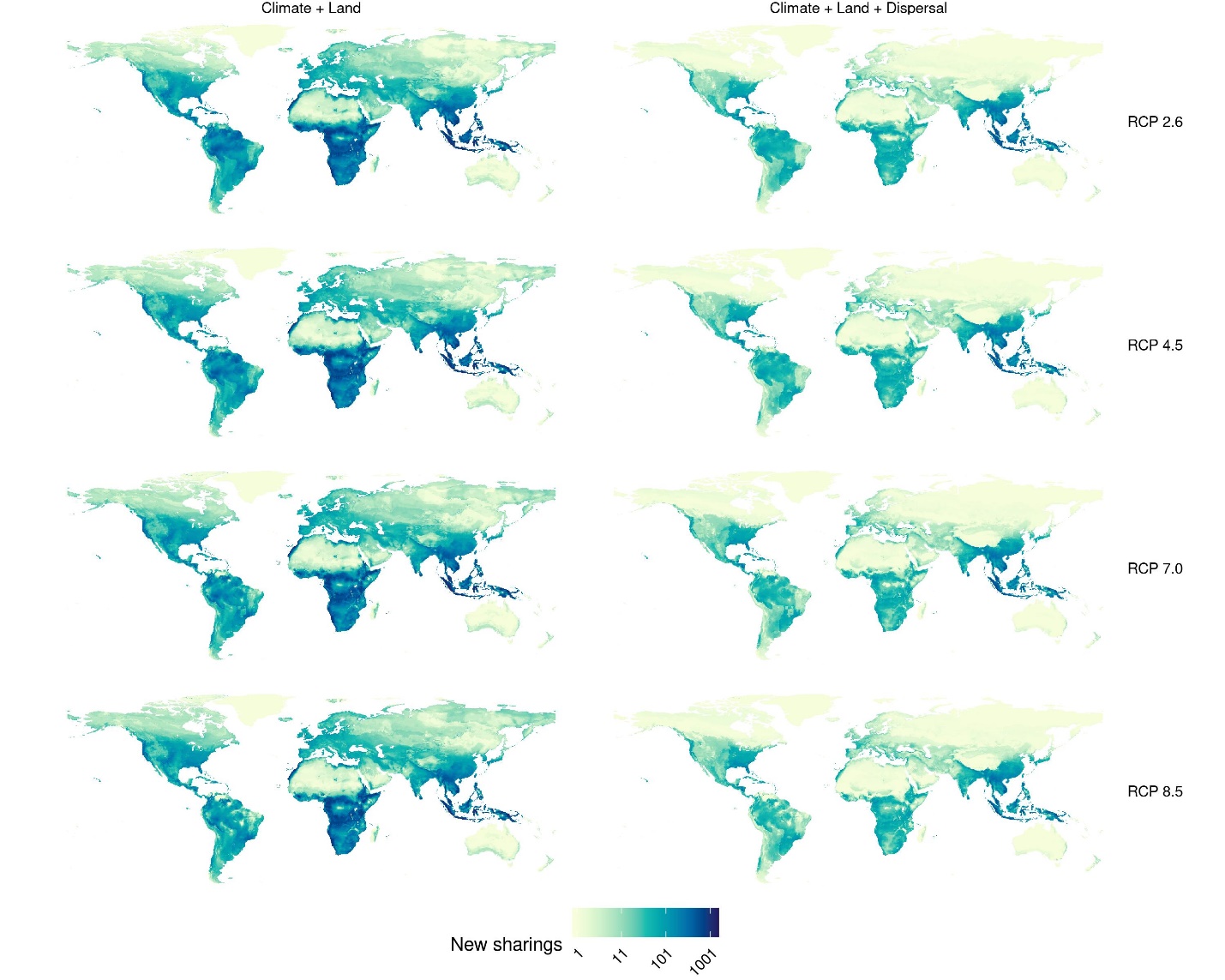

**Figure S18. Geographic distribution of viral sharing events in MRI-ESM2-0.** Predictions were carried out for four representative concentration pathways (RCPs), accounting for climate change and land use change, without (left) and with dispersal limits (right). Darker colours correspond to greater numbers of viral sharing events in the pixel.

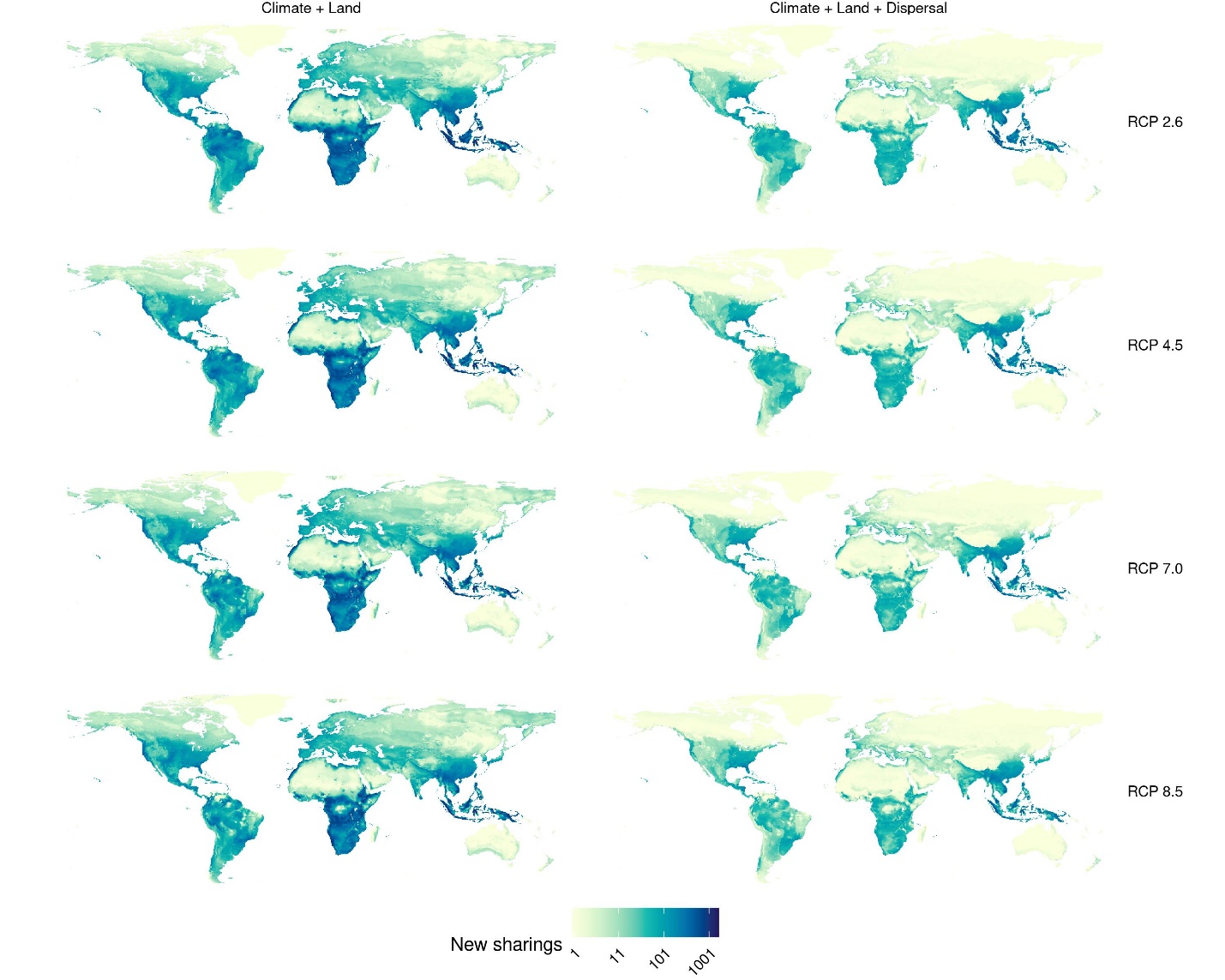
